# Systems Biomedicine of Rabies Delineates the Affected Signaling pathways

**DOI:** 10.1101/068817

**Authors:** Sadegh Azimzadeh Jamalkandi, Hamid Gholami Pourbadie, Mehdi Mirzaie, Sayed Hamid Reza Mozhgani, Farsheed Noorbakhsh, Behrouz Vaziri, Alireza Gholami, Naser Ansari-Pour, Mohieddin Jafari

**Author notes:** Corresponding Author: Mohieddin Jafari, Faculty Member, Pasteur Institute of Iran, 69, Pasteur St., 13164 Tehran, Iran. www.jafarilab-pasteur.com.

## Abstract

The prototypical neurotropic virus, rabies, is a member of the Rhabdoviridae family that causes lethal encephalomyelitis. Although there have been a plethora of studies investigating the etiological mechanism of the rabies virus and many precautionary methods have been implemented to avert the disease outbreak over the last century, the disease has surprisingly no definite remedy at its late stages. The psychological symptoms and the underlying etiology, as well as the rare survival rate from rabies encephalitis, has still remained a mystery. We, therefore, undertook a systems biomedicine approach to identify the network of gene products implicated in rabies. This was done by meta-analyzing whole-transcriptome microarray datasets of the CNS infected by strain CVS-11, and integrating them with interactome data using computational and statistical methods. We first determined the differentially expressed genes (DEGs) in each study and horizontally integrated the results at the mRNA and microRNA levels separately. A total of 61 seed genes involved in signal propagation system were obtained by means of unifying mRNA and microRNA detected integrated DEGs. We then reconstructed a refined protein-protein interaction network (PPIN) of infected cells to elucidate biological signaling transduction pathways affected by rabies virus. To validate our findings, we confirmed differential expression of randomly selected genes in the network using Real-time PCR. In conclusion, the identification of seed genes and their network neighborhood within the refined PPIN can be useful for demonstrating signaling pathways including interferon circumvent, toward proliferation and survival, and neuropathological clue, explaining the intricate underlying molecular neuropathology of rabies infection and thus rendered a molecular framework for predicting potential drug targets.

## Introduction

Growing evidence of inter-population and inter-individual variation in the attack rate and prognosis of specific infectious diseases suggest an underlying biological complexity. In fact, any perturbation in the densely organized inter-relationship of genetic and environmental factors, may lead to this intricate behavior (Hunter, 2005). In particular, the strange survival pattern observed from fatal rabies infection of the central nervous system (CNS) introduces this infection as a complex disease (De Souza and Madhusudana, 2014).

The prototypical neurotropic virus, rabies, is a member of the Rhabdoviridae family that causes lethal encephalomyelitis (Sugiura et al., 2011). The viruses in this family are enveloped with a single stranded negative sense RNA genome. The genomic length of the rabies virus (RABV) is about 12 kb and encodes five proteins including nucleoprotein (N), phosphoprotein (P), matrix protein (M), glycoprotein (G), and a viral RNA polymerase (L) (Yousaf et al., 2012). This neglected virus leads to death once the symptoms develop and has a mortality rate of 1:100,000 to 1:1,000 per year. The deceased intriguingly display no neural damage, neurohistopathological evidence or induced severe immune response (Schnell et al., 2010). In an organized hijacking program, the virus travels from the muscle tissue to the nervous system, migrates to the spinal cord and freely covers certain parts of the brain (Schnell et al., 2010). The virus spreads centrifugally to other organs and subsequently till next host. Although the host innate immune response including TLR, type 1 interferon, TNF alpha and IL-6 are the first defense line against a viral infection, this virus easily propagates in the nervous system. This suggests that the RABV has a specific mechanism to suppress host innate immunity (Rupprecht, 1996; Ito et al., 2011; Gomme et al., 2012b). Several laboratory strains of the RABV in addition to the wild types cause fatal acute encephalomyelitis associated with inflammation of the brain and spinal cord, leading to coma and death especially when the virus is injected intracerebrally in high dose (Meslin FX et al., 1996; Galelli et al., 2000; Baloul and Lafon, 2003; Baloul et al., 2004). In contrast to attenuated strains, wild type strains and CVS-11 do not induce histopathological changes indicative of apoptosis or necrosis in infected cells (Thoulouze et al., 1997; Lay et al., 2003; Prehaud et al., 2003; Thoulouze et al., 2003a; Thoulouze et al., 2003b). Accordingly, despite over 100 years of controlling rabies by developing RABV vaccines and serotherapy, the precise neurological and immunological etiology as well as rare survival cases from rabies encephalitis still remains a mystery (Gomme et al., 2012a; De Souza and Madhusudana, 2014).

After the emergence of omics technology, some studies have started to pave the way towards a better understanding of rabies fatal mechanism. Elucidating the essential biological processes involved in rabies progression has been based mainly on analyzing gene expression alterations (Jafari et al., 2013a). Zhao *et al*. reported expression profiling of mRNA and microRNA of rabies-infected cell (Zhao et al., 2011; Zhao et al., 2012a; Zhao et al., 2012b; Zhao et al., 2013). Suigiura *et al*. analyzed the gene expression profile of CNS tissue infected with CVS-11 (Sugiura et al., 2011). Changes in gene expression were also studied in marked neurons infected with recombinant RABV expressing CRE-recombinase (Gomme et al., 2012a). Numerous other studies have also analyzed gene expression profiling using transcriptomic or proteomic methods within diverse cellular models in different species (Wang et al., 2005; Dhingra et al., 2007; Fu et al., 2008; Zandi et al., 2009; Han et al., 2011; Thanomsridetchai et al., 2011; Wang et al., 2011; Vaziri et al., 2012; Farahtaj et al., 2013; Francischetti et al., 2013; Kluge et al., 2013; Silva et al., 2013; Venugopal et al., 2013; Zandi et al., 2013; Kasempimolporn et al., 2014; Kammouni et al., 2015; Mehta et al., 2015).

To increase the reliability of results and generalizability of these independent but related studies, it is recommended to statistically combine such data, commonly known as data integration or meta-analysis (Ramasamy et al., 2008). Several studies have shown the benefits of meta-analysis in terms of both higher statistical power and precision in detecting differentially expressed genes (DEGs) in different complex traits including infectious disease (Song et al., 2014; Camacho-Cáceres et al., 2015; Sharma et al., 2015; Wang et al., 2015a; Wang et al., 2015b; Yin et al., 2015). Further, data integration approaches at a higher level try to map multiple biological data levels into one mechanistic network to improve representativeness of data (Chen et al., 2008; Bowick and McAuley, 2011; Amiri et al., 2013; Depiereux et al., 2015; Paraboschi et al., 2015). The generated multi-dimensional network is likely to be more useful in inferring universally involved processes or pathways regardless of inter-studies differences (Azimzadeh et al., 2015).

Having in mind the common concerns in meta-analysis, we horizontally integrated some high-throughput transcriptome datasets to identify consensus DEGs. The underlying molecular network in rabies pathogenesis was then extracted based on protein-protein interaction network (PPIN) and signaling pathways by defining the identified DEGs as seed genes. Finally, using real-time PCR, we experimentally validated a number of key DEGs in rabies-infected cells. We demonstrate that a systems biomedicine approach, based on integrating omics datasets and experimental validation, may be used to shed light on a vague portrait of a complex disease pathobiology.

## Methods

### Super horizontal integration

#### Datasets

We looked into all databases pertaining to microarray data at both levels of mRNA and microRNA. This was done by searching databases (Gene Expression Omnibus (GEO), ArrayExpress, Google Scholar and PubMed NCBI) and studies regarding the rabies virus were extracted with rabies-related keywords including “rabies”, “RABV” and “rhabdoviridae” (Fig. 1A). Out of a total of 13 studies, nine were selected for further analysis. The list of included studies and their respective features are given in Supplementary File 1. In the majority of studies, “brain” and “brain spinal” were the tissues under investigation, and the rest were examined on *Mus musculus*-derived microglial cell line. It should also be noted that only 2 out of 9 inclusive studies were conducted on both mRNA and microRNA levels, with others analyzing only one level of data. The 4, 3 and 2 of these studies were performed on samples infected by CVS-11, FJDRV and ERA and RABV-Cre, respectively.

#### Data normalization

In order to prepare data for integration and detect DEGs, it is necessary to use preprocessed and background corrected microarray-data (Ramasamy et al., 2008). First, we checked the quality of recorded CELL format data all of which required to be normalized. The data were normalized by using the “Affy” package in Bioconductor (Gautier et al., 2004). This includes between and within array normalization which reduces the effect of noise and contributes to data consistency. The MA plot and qq-plot for the pairs of samples in each study was analyzed separately to check the normality of data after normalization as a quality control step. Although the qq-plot of some studies revealed that the normalization methods had worked fine, normalization was not successful in datasets with significantly small sample sizes mainly because normality assumptions are violated in low sample size studies.

#### Analysis of Differentially Expressed Genes (DEGs)

Assuming that microarray data are normally distributed, the routine procedure to detect DEGs is to perform t-test, however, this may result in misleading conclusions if the normality assumption is violated. A recent study showed that oligonucleotide expression values, resulting from widely acceptable calculation methods, are not normally distributed (Hardin and Wilson, 2009). This suggests that the results of t-test are biased and unreliable, especially when the sample size in each group is significantly small, and more robust methods should be implemented. Here, we used the Wilcoxon-Mann-Whitney nonparametric test as an alternative method to identify DEGs in each study with the significance level set to 0.05. The next step was to remove the unmapped probes and solve the problem of “many-to-many conversion” as described in (Ramasamy et al., 2008). This was done for both studies of mRNA and microRNA. The computational scripts plus all the raw data are provided in Supplementary file 2.

#### Meta-analysis

The results at each level (mRNA and microRNA) were integrated separately, using the inverse-variance technique and combining effect sizes as described in (Ramasamy et al., 2008). For integration purposes, the list of DEGs in each study was gathered and the effect size value of each gene was then calculated. We only selected genes with an absolute effect size value greater than 0.8 or those with a fold-change less than 0.33 or greater than 1.5 (in at least one study) as the frequently accepted cut-off for fold-change. We obtained two different lists of significant differentially expressed values for mRNA and microRNA, respectively. We obtained the target gene symbols of each microRNA accession ID using miRDB (http://mirdb.org) (Wong and Wang, 2015). To be best of our knowledge, for *Mus musculus*, this is the most up-to-date repository to convert accession IDs. Finally, after these two parallel horizontal integrations, the union and intersection of the results were extracted and the final gene list, namely the super-horizontally integrated DEGs (SHIDEGs), was generated.

### Background network construction

We used the STRING database to constructed a large-scale PPIN from seven available interaction sources and chose the lowest cut-off for combined scores (Downloaded on 2 September 2015) (Szklarczyk et al., 2014). A total of 8,604 proteins based on 9,162 SHIDEGs were represented in STRING. Accordingly, a total of 1,223,630 edges were extracted and the STRING combined scores were used as edge weights. Next, the weighted adjacency matrix was transformed to a new adjacency matrix using topological overlapping measure (TOM) function in WGCNA package of R software (Yip and Horvath, 2007; Langfelder and Horvath, 2008; Song et al., 2012). It should be noted that the TOM transformation increases the non-zero adjacency matrix elements as well as very low weight values in this case. The transformed weight distributions of STRING default cut-offs, from lowest to highest confidence, were thus considered to define a new threshold. The 3^rd^ quartile of transformed scores (0.4577) of the highest confidence was selected to strictly filter weak and false-positive interactions.

### Neighborhood ranking

Using the custom igraph package in R, we generated a matrix of all shortest paths between all pairs of nodes in a weighted network with the algorithm of Dijkstra (Csardi and Nepusz, 2006). First, we substituted raw weights with 1-weight to increase reachability of nodes with high weights to seed nodes in the shortest path finding procedure. We then defined a distance score, *D_j_*, for each node in the PPIN as the difference in average of the shortest path to the node when starting on a non–seed node compared with when starting on a seed node, normalized by the average shortest path to reach the node from the whole network.

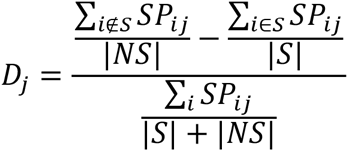

Here S is the set of nodes that fall into the seed gene set and NS is the set of nodes that are non-seed nodes. Therefore, a score greater than zero implies that node j falls closer on average to the seed nodes than it does on average to the rest of the network. The rabies network was generated based on the SHIDEGs seed list and each member of the seed list by scoring all nodes in the network and using a cutoff score of zero to define the neighborhood. It should be noted that the D scores were calculated without imposing any threshold on edge weights.

### Undirected PPIN; topological and pathway enrichment analysis

To reconstruct a high confidence PPIN around our seed gene set, we used the 0.4577 threshold to filter weak interaction among neighborhood nodes. This filtering resulted in the proximal neighborhood network of seeds (Table 3). Using Gephi version 0.9, the global topological properties of the resulting PPIN along with module identification was analyzed. To undertake enrichment analysis among the detected modules, ClueGO 2.1.7 (Bindea et al., 2009) in Cytoscape 3.2.1 was used based on *Mus musculus* using parameters update KEGG (Kanehisa et al., 2014), Reactome (Croft et al., 2014) and Wikipathway ontology databases (Kelder et al., 2012), default term selection options, hypergeometric test and Bonferroni step down p-value correction.

### Signaling network analysis

The rabies-implicated signaling network (RISN) was constructed based on the KEGG pathways enriched in the rabies PPIN. All statistically significant and frequent pathways in all PPIN modules were extracted and merged together to build a large-scale RISN. All SHIDEGs were then delineated in this network by different color labeling. After reviewing clinical and physiological evidences pertaining to the RABV, the whole RISN was delineated into a less complex network.

### Cell culture and virus

The Neuro-2a cell line, a murine neuroblastoma cell line, and CVS-11 strain of the RABV (the challenge virus standard) were obtained from the WHO collaborating center for reference and research on rabies, Pasteur Institute of Iran (Tehran, Iran). Virus titers were determined by a focal infectivity assay using BSR (a line of BHK) cells. Neuro-2a cells were grown in Dulbecco’s Modified Eagle Medium (DMEM) containing 4500 mg/L glucose and sodium bicarbonate, supplemented with 10% fetal bovine serum. Cultures were maintained at 37 C in a 5% CO2 humidified cell incubator with growth medium replaced every 48 hours. For all experiments, cells were subcultured into 25 cm tissue culture flasks and were grown for 16 hours before infection.

### Total RNA isolation, cDNA synthesis and primer design for PCR

Total mRNA was isolated from neuroblastoma cells (mock infected and infected with the CVS-11 strain of RABV) using the RNX RNA Isolation Kit (CinnaGen Inc., Tehran, Iran). The amount and purity of RNA were determined by Biotek microplate spectrophotometry. The extracted RNA was then treated with DNase to remove genomic DNA. Total RNA (1.7 μg/ml) was reverse transcribed into first-strand cDNA by the SuperScript III First-Strand Synthesis System (Thermo Fisher Scientific) and oligo(dT)18 according to the manufacturer’s protocol. Primer specificity was tested by primer-BLAST (http://blast.ncbi.nlm.nih.gov/Blast.cgi) and experimentally by the positive control amplification. Optimal PCR conditions were identified for each primer pair. GAPDH was used as an internal control for RT-polymerase chain reactions,.

### Quantitative real-time PCR

Reverse transcription-quantitative real-time PCR (RT-qPCR) was carried out on a Rotor-Gene Q 5plex HRM instrument (Qiagen, Hilden, Germany) with EvaGreen fluorescence dye (Biotium, Hayward, USA) to monitor cDNA amplification of *GNAI2*, *AKT3*, *IL21* and *GAPDH* through increased fluorescence intensity. The specificity of the amplified products was checked by melting curve analysis, and the expected size of the fragments was further visualized by gel electrophoresis (2% agarose) and staining with GelRed (Biotium, Hayward, CA). Results were confirmed by triplicate testing. Relative mRNA expression was calculated using the delta-delta Ct method (Livak and Schmittgen, 2001). Sequences were analyzed using Seqscanner. Statistical analysis was performed by depicting an error bar for each gene in each condition to compare relative expression of the abovementioned genes in uninfected and RABV-infected states.

## Results

This study comprises seven steps in two separate parts as illustrated in the Fig 1. After a systematic literature review, 9 transcriptomic datasets pertaining to rabies were collected (Supplementary File 1). DEGs were identified in the mRNA and microRNA datasets and used as seed genes to construct a PPIN of rabies infection. In the second part, analysis of the PPIN neighborhood and rabies-triggered signaling was implemented to create a mechanistic description of the molecular pathobiology of rabies infection (Fig. 1). These steps are discussed in more detail below.

**Fig 1.**
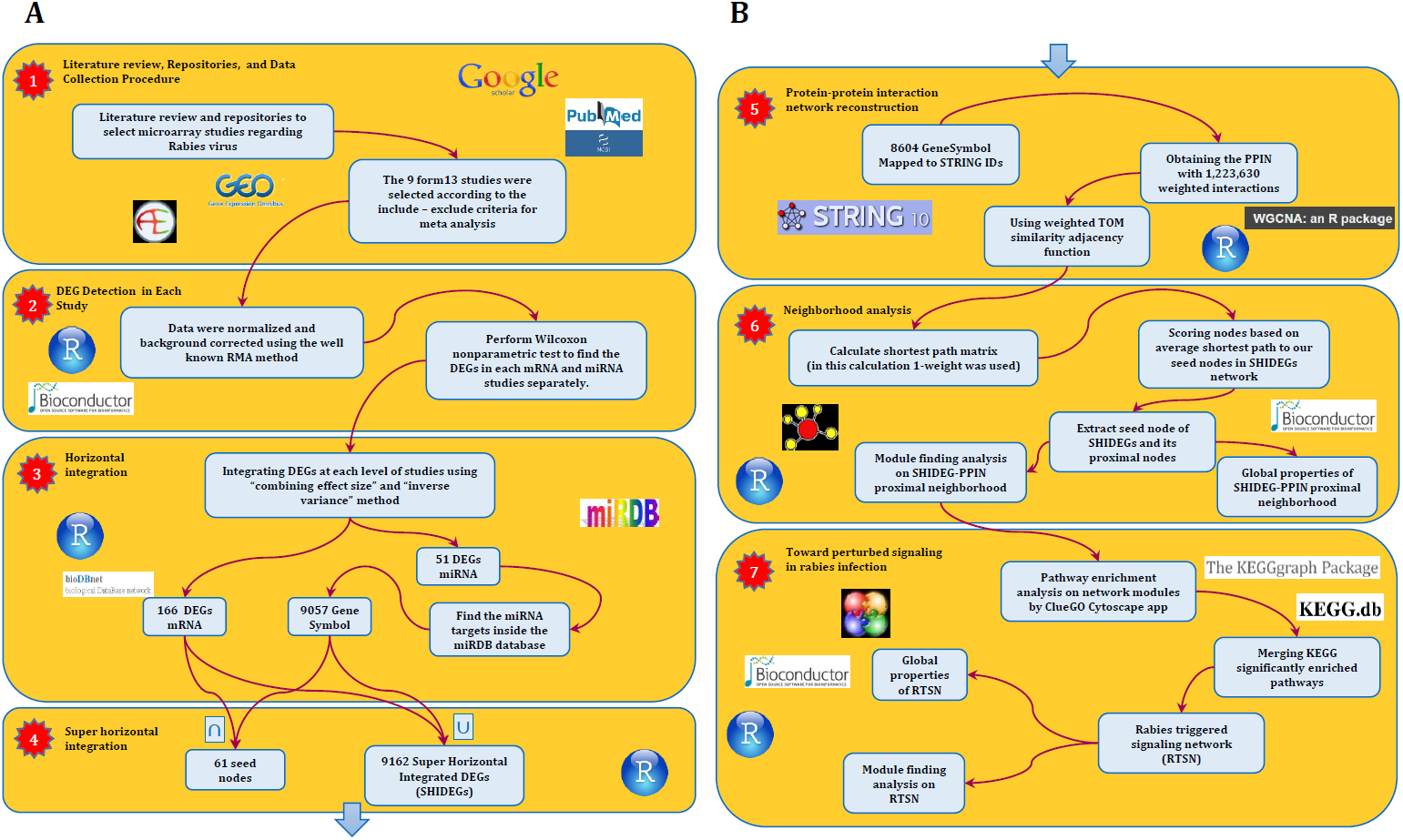
The abstract flowchart of this study design. A) The four steps taken to obtain the primitive interaction network of the RABV infection. The first step was a systematic review of the literature regarding the RABV. GEO was used to obtain the expression values of studies at the two levels of expression profiling by array of coding and non-coding RNA. Having selected the studies according to our data integration criteria, we moved on to the second step in which we detected DEGs at each level using nonparametric methods. The third step included integration of results using the meta-analysis techniques described in (Ramasamy et al., 2008). The implemented integration method revealed that 166 mRNA and 9057 microRNA are differentially expressed. The next step consisted of super horizontal integration of all transcriptome data. Finally, the 9162 expressed genes were mapped to STRING v 10.0 for further analysis. B)The next stage of the approach which comprised three steps resulted in the rabies-implicated signaling network (RISN). Firstly, the PPIN of all 9162 genes was reconstructed using STRING v10. The combined score calculated in STRING was used as edge weight in the SHIDEG-PPIN. In the second step, the proximal nodes of seed nodes in the whole SHIDEG-PPIN were found to create the seed neighborhood network. Module finding along was undertaken. Next, the significantly enriched signaling pathways were extracted in the network modules separately. These pathway were then merged together, forming the rabies-implicated signaling network (RISN). Finally, analysis the global network and functional module finding analysis was performed followed by biological inference.

### Union of mRNA and microRNA transcriptome data by super horizontal integration reveals an intriguing list of seed genes

A total of 166 DEGs were identified at the mRNA level. Analysis at the microRNA level led to the identification of 9057 genes. The genes at both mRNA and microRNA levels were combined to create a list of 9162 “super horizontally integrated – DEGs (SHIDEGs)” and a list of 61 intersecting genes considered as “seed genes” (Table 1). Of the 9162 SHIDEGs, 8604 (∼93%) were mapped to STRING version 10 and the obtained network contained 1,223,630 weighted protein-protein interactions.

**Table 1.**
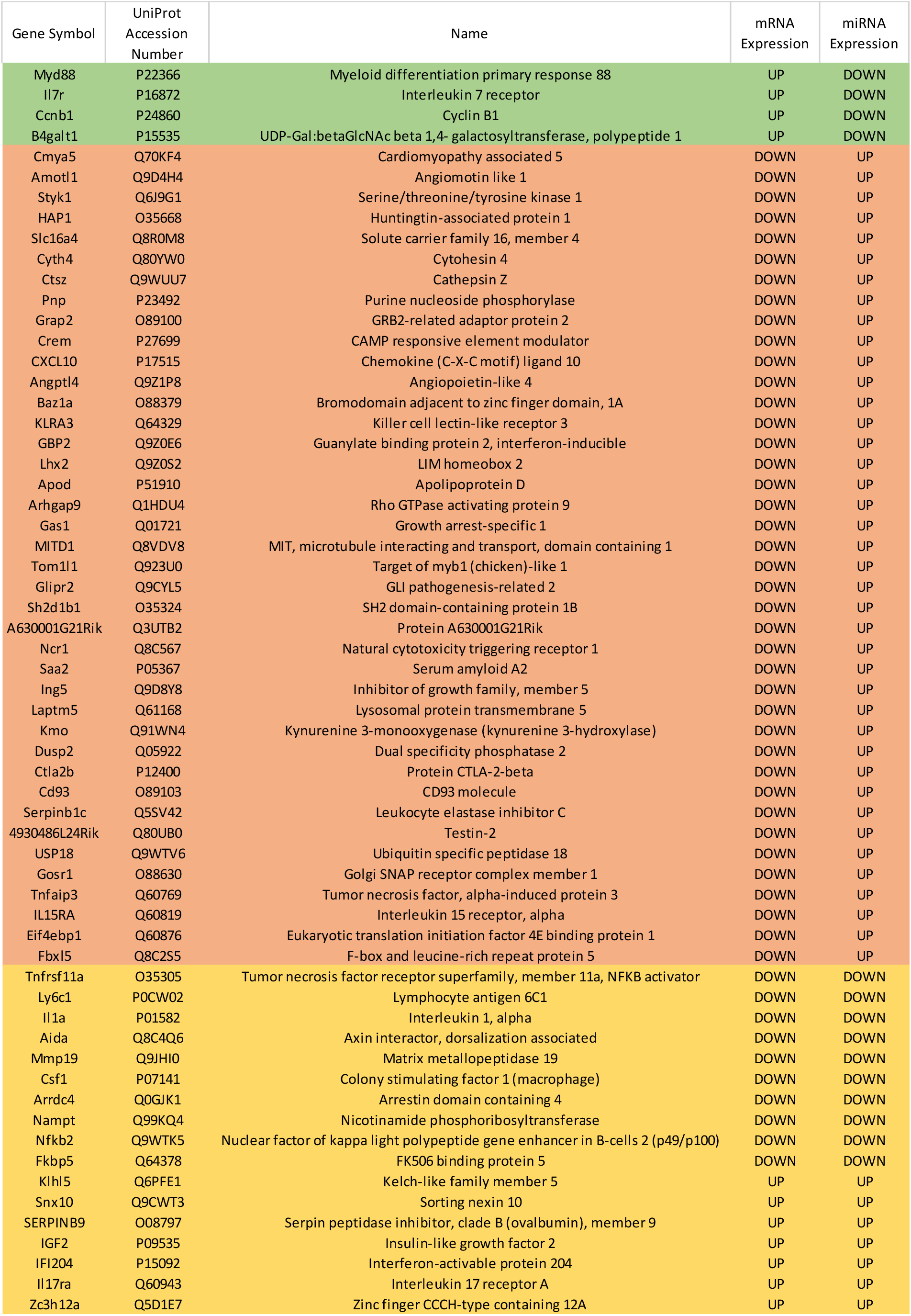
Seed genes. The 61 seed genes are presented in this table. The green, red and yellow color indicate over-, under and ambivalent expression of genes regarding mRNA and microRNA expression evidence.

| Gene Symbol | UniProt<br>Accession<br>Number | Name | mRNA<br>Expression | miRNA<br>Expression |
| --- | --- | --- | --- | --- |
| Myd88 | P22366 | Myeloid differentiation primary response 88 | UP | DOWN |
| Il7r | P16872 | Interleukin 7 receptor | UP | DOWN |
| Ccnb1 | P24860 | Cyclin B1 | UP | DOWN |
| B4galt1 | P15535 | UDP-Gal:betaGlcNAc beta 1,4- galactosyltransferase, polypeptide 1 | UP | DOWN |
| Cmya5 | Q70KF4 | Cardiomyopathy associated 5 | DOWN | UP |
| Amotl1 | Q9D4H4 | Angiomotin like 1 | DOWN | UP |
| Styk1 | Q6J9G1 | Serine/threonine/tyrosine kinase 1 | DOWN | UP |
| HAP1 | O35668 | Huntingtin-associated protein 1 | DOWN | UP |
| Slc16a4 | Q8R0M8 | Solute carrier family 16, member 4 | DOWN | UP |
| Cyth4 | Q80YW0 | Cytohesin 4 | DOWN | UP |
| Ctsz | Q9WUU7 | Cathepsin Z | DOWN | UP |
| Pnp | P23492 | Purine nucleoside phosphorylase | DOWN | UP |
| Grap2 | O89100 | GRB2-related adaptor protein 2 | DOWN | UP |
| Crem | P27699 | CAMP responsive element modulator | DOWN | UP |
| CXCL10 | P17515 | Chemokine (C-X-C motif) ligand 10 | DOWN | UP |
| Angptl4 | Q9Z1P8 | Angiopietin-like 4 | DOWN | UP |
| Baz1a | O88379 | Bromodomain adjacent to zinc finger domain, 1A | DOWN | UP |
| KLRA3 | Q64329 | Killer cell lectin-like receptor 3 | DOWN | UP |
| GBP2 | Q9Z0E6 | Guanylate binding protein 2, interferon-inducible | DOWN | UP |
| Lhx2 | Q9Z0S2 | LIM homeobox 2 | DOWN | UP |
| Apod | P51910 | Apolipoprotein D | DOWN | UP |
| Arhgap9 | Q1HDU4 | Rho GTPase activating protein 9 | DOWN | UP |
| Gas1 | Q01721 | Growth arrest-specific 1 | DOWN | UP |
| MITD1 | Q8VDV8 | MIT, microtubule interacting and transport, domain containing 1 | DOWN | UP |
| Tom1l1 | Q923U0 | Target of myb1 (chicken)-like 1 | DOWN | UP |
| Glipr2 | Q9CYL5 | GLI pathogenesis-related 2 | DOWN | UP |
| Sh2d1b1 | O35324 | SH2 domain-containing protein 1B | DOWN | UP |
| A630001G21Rik | Q3UTB2 | Protein A630001G21Rik | DOWN | UP |
| Ncr1 | Q8C567 | Natural cytotoxicity triggering receptor 1 | DOWN | UP |
| Saa2 | P05367 | Serum amyloid A2 | DOWN | UP |
| Ing5 | Q9D8Y8 | Inhibitor of growth family, member 5 | DOWN | UP |
| Laptm5 | Q61168 | Lysosomal protein transmembrane 5 | DOWN | UP |
| Kmo | Q91WN4 | Kynurenine 3-monooxygenase (kynurenine 3-hydroxylase) | DOWN | UP |
| Dusp2 | Q05922 | Dual specificity phosphatase 2 | DOWN | UP |
| Ctla2b | P12400 | Protein CTLA-2-beta | DOWN | UP |
| Cd93 | O89103 | CD93 molecule | DOWN | UP |
| Serpinb1c | Q5SV42 | Leukocyte elastase inhibitor C | DOWN | UP |
| 4930486L24Rik | Q80UB0 | Testin-2 | DOWN | UP |
| USP18 | Q9WTV6 | Ubiquitin specific peptidase 18 | DOWN | UP |
| Gosr1 | O88630 | Golgi SNAP receptor complex member 1 | DOWN | UP |
| Tnfaip3 | Q60769 | Tumor necrosis factor, alpha-induced protein 3 | DOWN | UP |
| IL15RA | Q60819 | Interleukin 15 receptor, alpha | DOWN | UP |
| Eif4ebp1 | Q60876 | Eukaryotic translation initiation factor 4E binding protein 1 | DOWN | UP |
| Fbxl5 | Q8C2S5 | F-box and leucine-rich repeat protein 5 | DOWN | UP |
| Tnfrsf11a | O35305 | Tumor necrosis factor receptor superfamily, member 11a, NFKB activator | DOWN | DOWN |
| Ly6c1 | P0CW02 | Lymphocyte antigen 6C1 | DOWN | DOWN |
| Il1a | P01582 | Interleukin 1, alpha | DOWN | DOWN |
| Aida | Q8C4Q6 | Axin interactor, dorsalization associated | DOWN | DOWN |
| Mmp19 | Q9JHI0 | Matrix metalloproteinase 19 | DOWN | DOWN |
| Csf1 | P07141 | Colony stimulating factor 1 (macrophage) | DOWN | DOWN |
| Arrdc4 | Q0GJK1 | Arrestin domain containing 4 | DOWN | DOWN |
| Nampt | Q99KQ4 | Nicotinamide phosphoribosyltransferase | DOWN | DOWN |
| Nfkb2 | Q9WTK5 | Nuclear factor of kappa light polypeptide gene enhancer in B-cells 2 (p49/p100) | DOWN | DOWN |
| Fkbp5 | Q64378 | FK506 binding protein 5 | DOWN | DOWN |
| Klhl5 | Q6PFE1 | Kelch-like family member 5 | UP | UP |
| Snx10 | Q9CWT3 | Sorting nexin 10 | UP | UP |
| SERPINB9 | O08797 | Serpin peptidase inhibitor, clade B (ovalbumin), member 9 | UP | UP |
| IGF2 | P09535 | Insulin-like growth factor 2 | UP | UP |
| IFI204 | P15092 | Interferon-activable protein 204 | UP | UP |
| Il17ra | Q60943 | Interleukin 17 receptor A | UP | UP |
| Zc3h12a | Q5D1E7 | Zinc finger CCCH-type containing 12A | UP | UP |

Next, we obtained gene ontology (GO) classifications for all seed genes. Using the Enrichr web based tools, significantly enriched biological process (BP), molecular function (MF) and cellular component (CC) terms were retrieved and then ranked by combined scores (Fig. 2). From a biological process point of view, diverse inflammatory responses such as cytokine-mediated signaling pathway (GO:0019221) and regulation of leukocyte activation (GO:0002694) were enriched. Also, the other moiety of BPs was generally associated with nucleotide biosynthetic processes (Fig. 2A). This observation confirmed the role of immune signaling pathways and propagation apparatus in rabies infection. CC and MF enriched terms further accentuated the role of signaling alteration in development of rabies (Fig. 2B-C).

**Fig 2.**
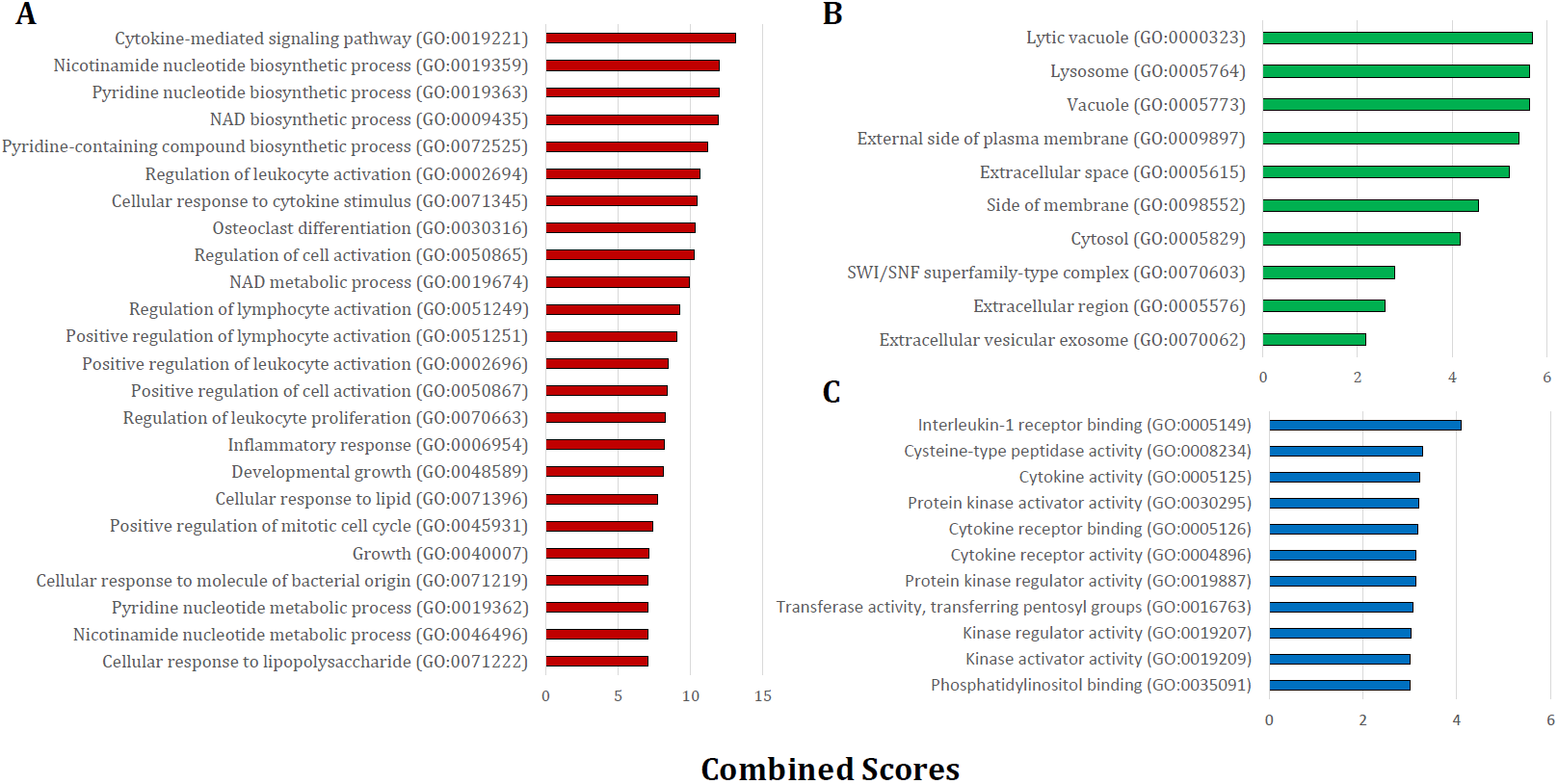
The significant GO terms based on the seed gene set. The enriched GO terms within (A) biological processes, (B) cellular components and (C) molecular functions are presented separately. All terms were statistically significant (P < 0.05) and are ranked based on Enrichr combined scores.

### Shortest path-based scoring allows identification of seed gene neighborhood in the protein-protein interaction space

We applied network concepts to explore more thoroughly the potential functional relationship between the identified DEGs and RABVetiology. We postulated that all integrated DEGs are involved in the global interactome perturbed by the RABV. We assume that the SHIDEGs are more confirmed to interact directly with the RABV and the neighbors of SHIDEGs are of the next level of etiological importance. To identify the disease subnetwork of SHIDEGs in PPIN, we retrieved the entire protein-protein weighted interactions from STRING (Szklarczyk et al., 2014). The giant component comprising 8604 DEG products was selected for further analysis. Given the high false-positive rate in PPINs (Jafari et al., 2013b; Jafari et al., 2015), the topological overlap matrix (TOM)-based adjacency function was used to filter the effect of spurious or weak connections (Li and Horvath, 2007; Yip and Horvath, 2007). Proteins encoded by the 61 seed genes were identified in the refined global PPIN for neighborhood analysis.

Based on biological parsimony and the observed patterns in different signaling databases, biological responses are controlled via a short signaling cascade (Gitter et al., 2011; Silverbush and Sharan, 2014). We therefore used the shortest path algorithm to identify nodes in proximity of the seed nodes. We then ranked the nodes within the whole PPIN using distance D. From the total 8,602 nodes within the robust PPIN, 3,775 nodes had a positive score and therefore fell within the seed gene neighborhood.

Subsequently, to filter the edges having low weight, STRING combined score (weight of edges) were transformed using the TOM-based adjacency function and those above the 3^rd^ quartile were retained. Of all nodes with D>0, 694 nodes passed this filter and were considered as the “seed gene proximal neighborhood network”. This resulted in the selection of highly important relationships among nodes based on network topology (Fig 3).

**Fig 3.**
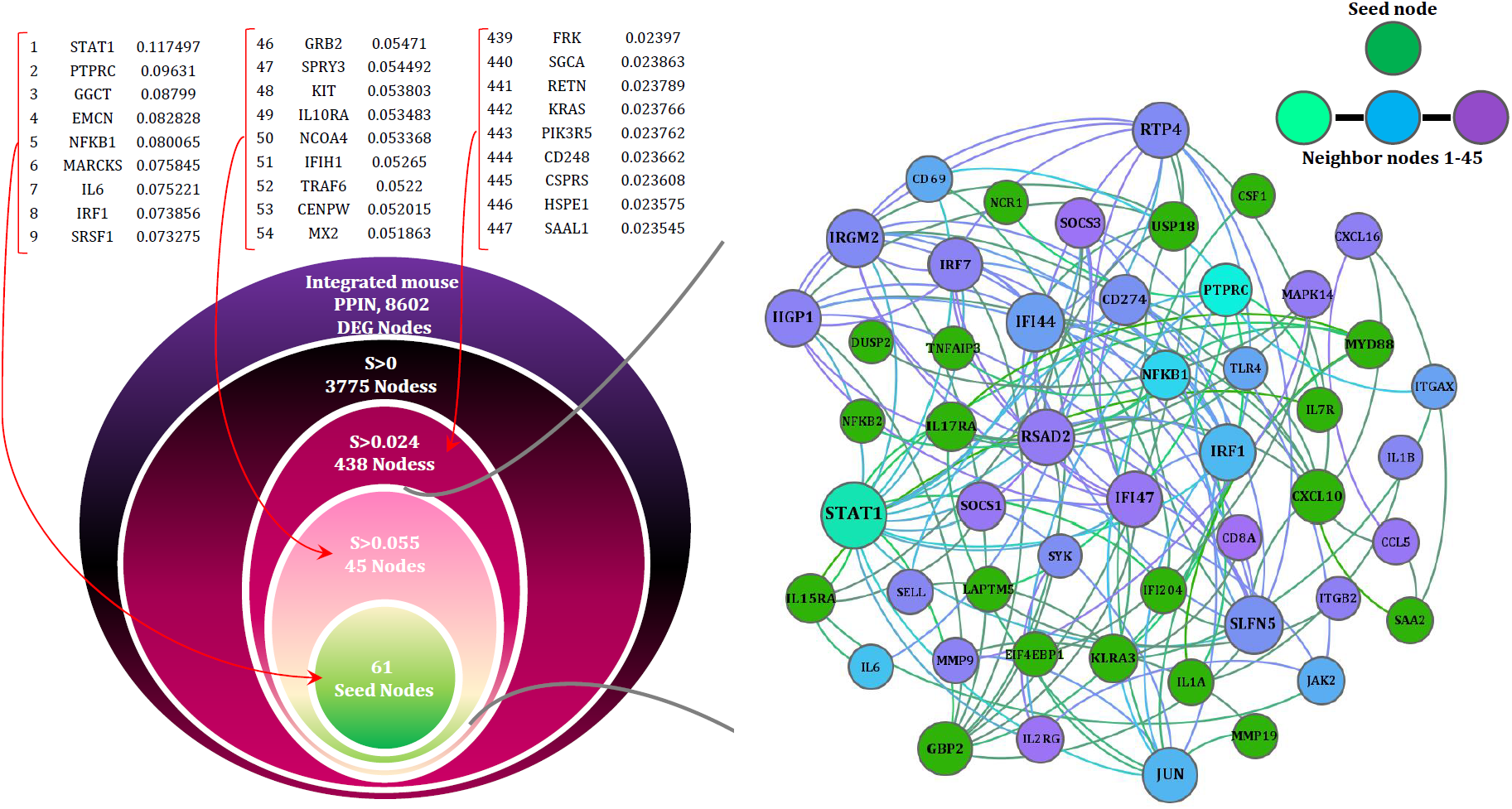
The network (SHIDEG-PPIN) onion diagram. A) Identification of the rabies disease neighborhood network based on proximal nodes of seed genes. The PPIN of all DEG products (8602/9162) existed in STRING were ranked on the basis of their shortest path (SP) score with the 61 rabies SHIDEGs as seed nodes. Selecting various score cutoffs (99.5th, 95th and 56th quantiles) allow neighborhoods of various sizes to be defined as shown in the nested circles. The ranked list of neighbors of seed nodes demonstrated with their scores. B) The minimized PPIN formed with the confident interactions (0.4 cut-off selected based on the TOM procedure) between the 61 seed nodes (green) and top 45 ranked proximal nodes (range of colors from cyan to purple) is shown at the top with node size representing degree of nodes.

### The identification of rabies-implicated gene products

The final rabies infection PPIN contained 694 nodes with 6097 interactions. The degree distribution (Supplementary File 3) and modularity index (∼0.7) of this PPIN indicate that it has a modular structure and a scale free topology. Its average path length and diameter were 5.16 and 14, respectively, showing that this relatively large and sparse network is small-world. To infer the functionality of this refined network, we analyzed the network modules. Twelve modules were detected by the fast unfolding clustering algorithm implemented in Gephi (V. 0.9) (Bastian et al.).

To avoid bias in inferring global properties of the network, the top three central nodes in each module were specifically shown in Fig 4. This figure illustrates these nodes in terms of degree and betweenness centrality. Interestingly, all of these nodes have diverse receptor binding and kinase activity functions based on GO enrichment analysis. On the other hand, our results revealed that inter-modular high-degree nodes related to CCR chemokine receptor binding (GO:0048020), R-SMAD binding (GO:0070412), responses to mechanical stimulus (GO:0009612), JAK-STAT cascade involved in growth hormone signaling pathway (GO:0060397) and negative regulation of neuron death (GO:1901215) were down-regulated by the RABV. In contrast, the local and global hub proteins associated with positive regulation of protein kinase activity (GO:0045860), cellular response to lipid (GO:0071396), neurotrophin TRK receptor signaling pathway (GO:0048011), G-protein coupled receptor binding (GO:0001664) and neuropeptide hormone activity (GO:0005184) were up-regulated, thus facilitating virus survival and propagation by avoiding programmed cell death.

**Fig 4.**
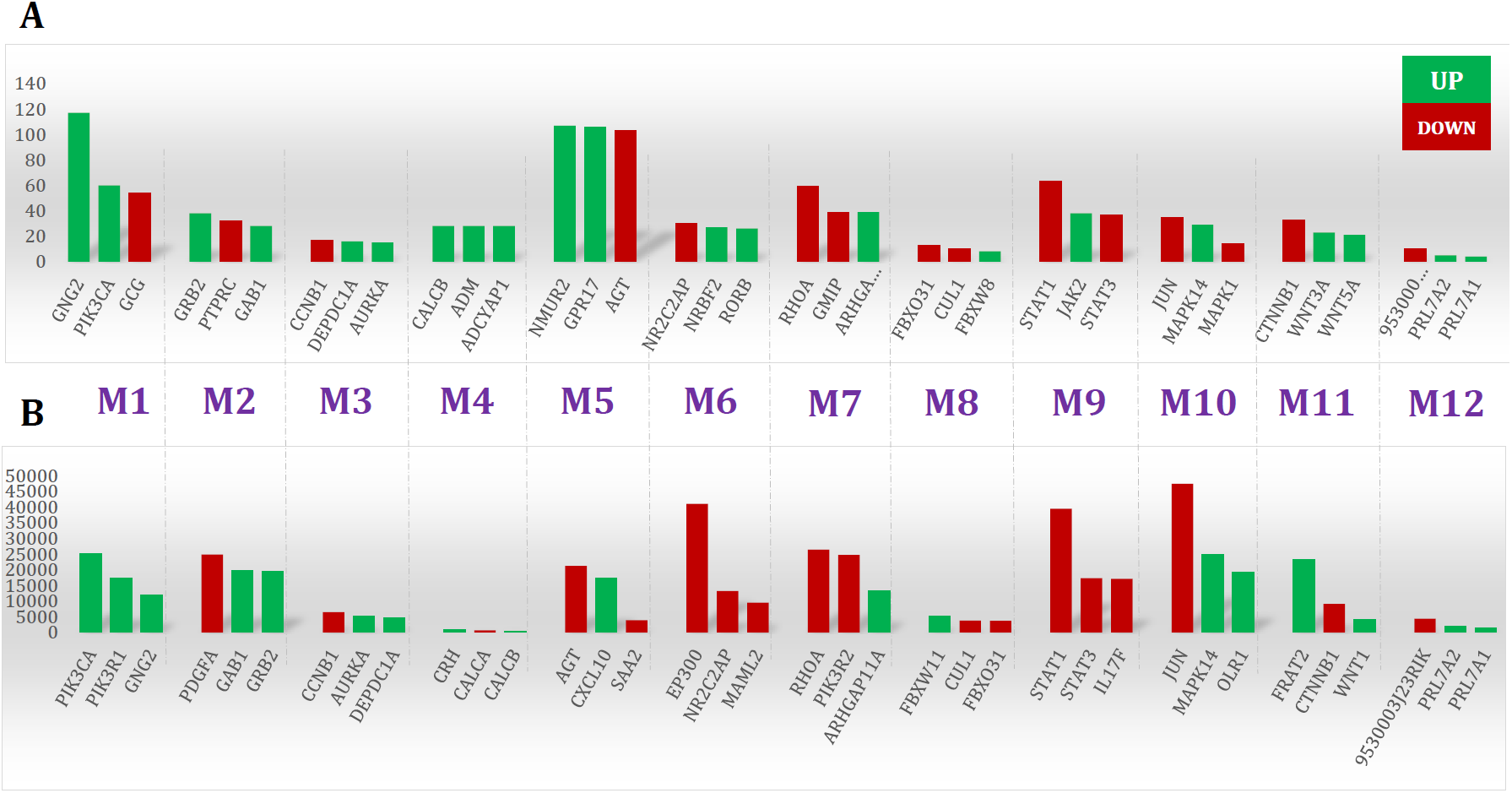
Top ranked nodes based on two centrality measures. The (A) degree and (B) betweenness centrality were calculated in the proximal neighborhood network to seed genes.

The top three ranked nodes of the 12 modules are presented separately in Fig 5. The over-expression and under-expression of these gene products are labeled by color. The same scenario also applied to nodes with high betweenness centrality. For example, nodes associated with neurotrophin receptor binding (GO:0005165) and cellular response to organonitrogen compound (GO:0071417) were over-expressed while those associated with natural immune system were underexpressed (Supplementary File 4). On top of that, the top five ranked nodes based on betweenness centrality, namely EP300, STAT1, RHOA, and PDGFA were under-expressed concurrently. This may lead to a lack of network coordination among different immune processes.

**Fig 5.**
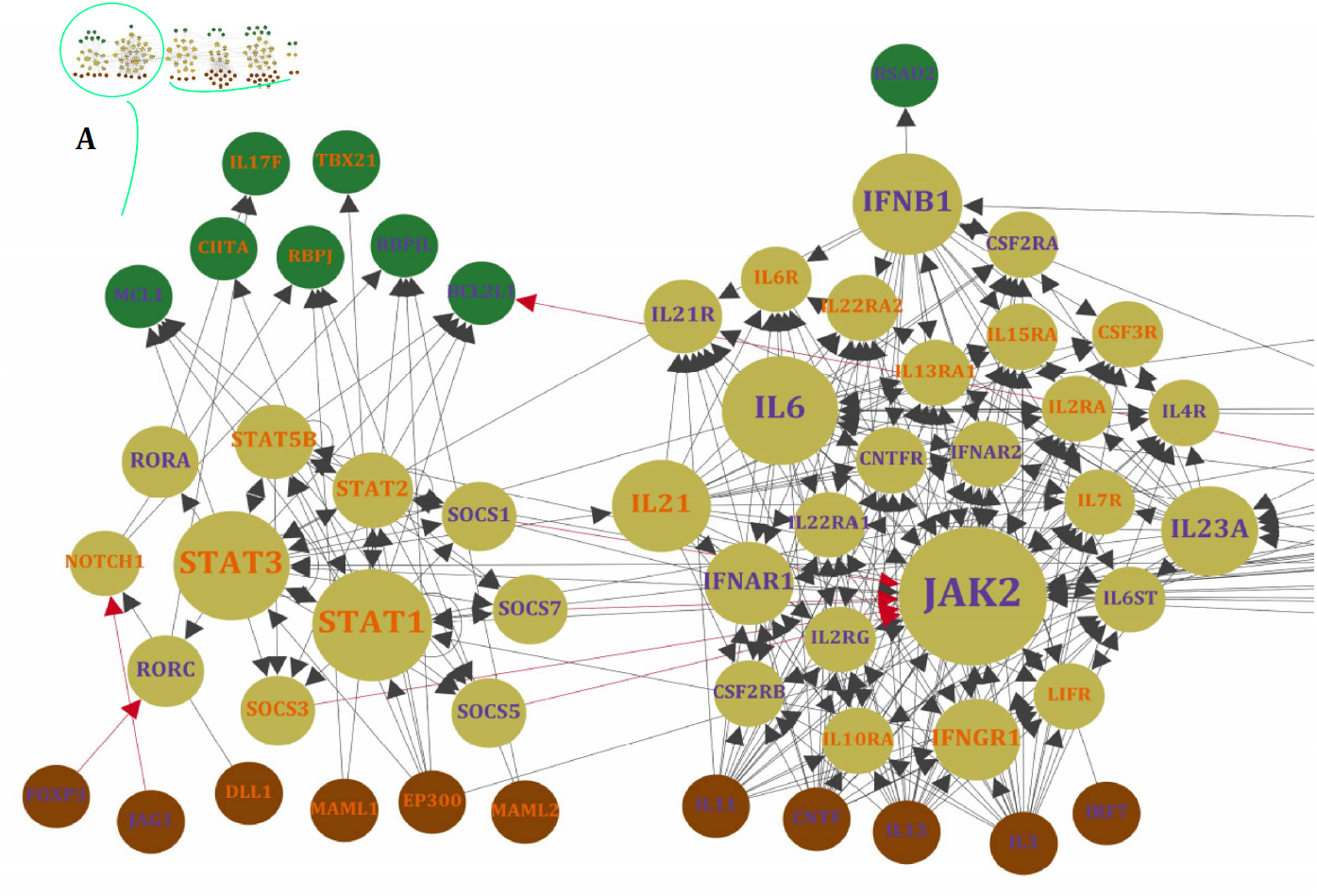
A bird’s eye view of RISN. The 22 over-expressed genes of significantly enriched KEGG pathways were selected and merged. The DEGs are labeled by different colors including orange and blue which indicates down-regulation and up-regulation of genes, respectively. The activator/inhibitor edges are also colored differently (red edges are inhibitors and black vice versa). The node colors represent its relative position in a directed network from brown (source nodes), cream (internal nodes) to green (sink nodes) and the node size is proportional to betweenness centrality value. In addition,, the six detected modules in three parts are displayed separately: (A) Interferon circumvent, (B) Toward proliferation and survival and (C) neuropathological clue. In this Figure, the six independent modules are represented in three colors; brown, cream and green, indicated root, internal and leaf node in this directed graph. Node size represents betweenness centrality of nodes and the red edges shows inhibitory interactions. To summarize the network for visualization, the SHIDEG nodes are only represented in this Figure.

Given the abundance of receptors and kinases in this network, we performed pathway enrichment analysis on each module separately. Using ClueGO (Cytoscape plugin (Bindea et al., 2009)), the statistically significant pathway terms were identified among those in the Kyoto Encyclopedia of Genes and Genomes (KEGG) (Kanehisa et al., 2014) and Reactome (Croft et al., 2014) databases (Table 2 and Supplementary File 5). We then ranked the enriched pathway terms based on gene coverage (Ansari-Pour et al., 2016).

**Table 2.** The enriched KEGG and Reactome pathways. The statistically significantly enriched KEGG and Reactome pathways were identified by ClueGO. The top three representative pathways identified in each refined SHIDEG-PPIN module are given along with their corrected p-values. The highlighted pathway names were found to be enriched in more than one module.

| Module No. | KEGG | Frequency | PValue Corrected with Bonferroni step down | Reactome | Frequency | PValue Corrected with Bonferroni step down |
| --- | --- | --- | --- | --- | --- | --- |
| M1 | vasopressin-regulated water reabsorption | 26 | 1.13E-33 | Peptide ligand-binding receptors | 22 | 9.62E-30 |
|  | Calcium signaling pathway | 21 | 2.76E-28 | Platelet activation, signaling and aggregation | 10 | 2.76E-08 |
|  | cGMP-PKG signaling pathway | 7 | 2.56E-05 | Thrombin signalling through proteinase activated receptors (PARs) | 7 | 2.49E-10 |
| M2 | PI3K-Akt signaling pathway | 30 | 2.16E-24 | Immune System | 49 | 9.78E-28 |
|  | Pathways in cancer | 30 | 1.26E-22 | Innate Immune System | 38 | 1.31E-25 |
|  | Ras signaling pathway | 29 | 2.10E-28 | Adaptive Immune System | 35 | 6.87E-23 |
| M3 | Cell cycle | 10 | 1.71E-11 | Cell Cycle | 28 | 4.52E-31 |
|  | Vasopressin-regulated water reabsorption | 2 | 2.42E-02 | Cell Cycle, Mitotic | 27 | 4.64E-31 |
|  | Bladder cancer | 2 | 4.17E-02 | M Phase | 12 | 1.80E-10 |
| M4 | Neuroactive ligand-receptor interaction | 16 | 3.08E-22 | G alpha (s) signalling events | 23 | 1.04E-47 |
|  | - | - | - | GPCR ligand binding | 21 | 1.91E-29 |
|  | - | - | - | Class B/2 (Secretin family receptors) | 10 | 5.73E-16 |
| M5 | Chemokine signaling pathway | 25 | 1.78E-29 | Signaling by GPCR | 68 | 2.40E-78 |
|  | Neuroactive ligand-receptor interaction | 25 | 2.85E-25 | GPCR downstream signaling | 63 | 2.90E-70 |
|  | Cytokine-cytokine receptor interaction | 20 | 4.75E-18 | G alpha (i) signalling events | 63 | 1.24E-110 |
| M6 | Thyroid hormone signaling pathway | 12 | 1.56E-12 | Generic Transcription Pathway | 46 | 3.10E-54 |
|  | Notch signaling pathway | 8 | 8.68E-10 | Developmental Biology | 28 | 1.47E-24 |
|  | Maturity onset diabetes of the young | 3 | 7.71E-03 | Nuclear Receptor transcription pathway | 27 | 4.31E-53 |
| M7 | Regulation of actin cytoskeleton | 9 | 1.74E-06 | Signaling by Rho GTPases | 52 | 1.49E-77 |
|  | Pancreatic cancer | 3 | 3.45E-02 | Rho GTPase cycle | 51 | 1.99E-104 |
|  | - | - | - | G alpha (12/13) signalling events | 17 | 1.45E-25 |
| M8 | Ubiquitin mediated proteolysis | 8 | 2.36E-12 | Association of Tric/CCT with target proteins during biosynthesis | 4 | 1.40E-07 |
|  | Circadian rhythm | 2 | 1.19E-03 | Protein folding | 4 | 1.44E-06 |
|  | - | - | - | Chaperonin-mediated protein folding | 4 | 1.04E-06 |
| M9 | Jak-STAT signaling pathway | 37 | 2.06E-50 | Immune System | 57 | 3.06E-41 |
|  | Cytokine-cytokine receptor interaction | 36 | 1.80E-39 | Cytokine Signaling in Immune system | 53 | 1.54E-67 |
|  | Measles | 26 | 9.44E-32 | Interferon Signaling | 31 | 1.08E-35 |
| M10 | MAPK signaling pathway | 25 | 3.59E-24 | Innate Immune System | 31 | 2.54E-21 |
|  | Pathways in cancer | 23 | 7.36E-17 | Toll-Like Receptors Cascades | 19 | 1.54E-20 |
|  | PI3K-Akt signaling pathway | 22 | 7.01E-17 | Toll Like Receptor 3 (TLR3) Cascade | 18 | 7.97E-22 |
| M11 | Wnt signaling pathway | 33 | 6.80E-57 | Signaling by Wnt | 30 | 1.06E-37 |
|  | Pathways in cancer | 28 | 2.05E-31 | TCF dependent signaling in response to WNT | 23 | 1.72E-28 |
|  | Melanogenesis | 26 | 2.21E-44 | Class B/2 (Secretin family receptors) | 18 | 1.63E-27 |
| M12 | - | - | - | Amyloids | 5 | 1.21E-09 |
|  | - | - | - | Disease | 5 | 8.05E-05 |
|  | - | - | - |  |  |  |

Furthermore, to evaluate the quality of module discovery results, conformity of enriched pathways in a module was assessed with respect to the interconnectedness level of that module (Supplementary File 6). Our results demonstrated that the KEGG enriched pathway similarity matrix was significantly correlated with the module interconnectivity matrix (P-value < 0.01) and that they were highly similar (Rand measure = 73%).

### Towards identifying the signaling network involved in rabies etiology

In order to retrieve casual relationships, we used the KEGG database and enrichment results to prune the proximal network of seed genes. By reviewing the significantly enriched pathways, all KEGG pathways (N=47; Supplementary File 5) were merged to reconstruct the enriched signaling network pertaining to rabies etiology. The full signaling network is presented in Supplementary File 7, but the merge of only 22 of them were presented in Fig 5. These pathways were selected based on gene coverage, reported relevance to rabies pathobiology and association with other viral infections. Then, the DEGs related to these pathways were used to mine the rabies-implicated signaling network (RISN) based on the following KEGG pathways: PI3K-AKT signaling pathway (KEGG:04151), cell cycle (KEGG:04110), Jak-STAT signaling pathway (KEGG:04630), circadian rhythm (KEGG:04710), pertussis (KEGG:05133), leishmaniasis (KEGG:05140), tuberculosis (KEGG:05152), hepatitis B (KEGG:05161), influenza A (KEGG:05164), herpes simplex infection (KEGG:05168), Epstein-Barr virus infection (KEGG:05169), inflammatory bowel disease (IBD) (KEGG:05321), PPAR signaling pathway (KEGG:03320), hematopoietic cell lineage (KEGG:04640), neuroactive ligand-receptor interaction (KEGG:04080), Notch signaling pathway (KEGG:04330), inflammatory mediator regulation of TRP channels (KEGG:04750), TNF signaling pathway (KEGG:04668), T cell receptor signaling pathway (KEGG:04660), cytokine-cytokine receptor interaction (KEGG:04060), chemokine signaling pathway (KEGG:04062) and ubiquitin mediated proteolysis (KEGG:04120). The DEGs are showed by different label colors. Downregulated and upregulated genes are labeled as orange and blue, respectively. The activator/inhibitor edges are also colored differently (Red edges are inhibitor and black vice versa). The nodes’ color represent the position of them in a directed network from brown (Source nodes), cream (Internal nodes) to green (Sink nodes) and the node size is proportional to betweenness value. Also, the six detected modules in the network are displayed separately. The important sink and source nodes along with the nodes with high betweenness centrality in the whole RISN are listed in Table 3. The influence of nodes with high betweenness on propagating or focusing information among this signaling network is presented by the information release index (IRI), *IRI* = log (*Outdegree/Indegree*). Next, we analyzed RISN in the following categories, based on biological relevance.

**Table 3.** Details of important sink and source nodes along with nodes with top 26 betweenness centrality values in the whole RISN.

| Gene symbol | Protein name | Po. | U/D | Role in rabies |
| --- | --- | --- | --- | --- |
| **CCL11** | Chemokine (C-C motif) ligand 11 | Source | Down |  |
| **CCL20** | Chemokine (C-C motif) ligand 20 | Source | Down |  |
| **CCL22** | Chemokine (C-C motif) ligand 22 | Source | Up |  |
| **CCL28** | Chemokine (C-C motif) ligand 28 | Source | Up |  |
| **CCL3** | Chemokine (C-C motif) ligand 3 | Source | Up | Highly activated in the brains of mice infective with Rabies (71) |
| **CCL5** | Chemokine (C-C motif) ligand 5 | Source | Up | Known as a vital regulator which is involved in convincing encephalomyelitis (72), raised levels of mRNA transcripts (73), the expression value of CXCL10 and CCL5 in microglia is accurately regulated while the multiple signaling pathways are activated. |
| **CD28** | CD28 molecule | Source | Up |  |
| **CDC37** | Cell division cycle 37 | Source | Up |  |
| **CNTF** | Ciliary neurotrophic factor | Source | Up |  |
| **CX3CL1** | Chemokine (C-X3-C motif) ligand 1 | Source | Up |  |
| **CXCL1** | Chemokine (C-X-C motif) ligand 1 (melanoma growth stimulating activity, alpha) | Source | Up |  |
| **CXCL10** | Chemokine (C-X-C motif) ligand 10 | Source | Up | Known as a vital regulator which is involved in convincing encephalomyelitis (72), raised levels of mRNA transcripts (73), the expression value of CXCL10 and CCL5 in microglia is accurately regulated while the multiple signaling pathways are activated. |
| **CXCL12** | Chemokine (C-X-C motif) ligand 12 | Source | Up |  |
| **CXCL16** | Chemokine (C-X-C motif) ligand 16 | Source | Down |  |
| **CXCL2** | Chemokine (C-X-C motif) ligand 2 | Source | Up |  |
| **CXCL5** | Chemokine (C-X-C motif) ligand 5 | Source | Up |  |
| **DLL1** | Delta-like 1 (Drosophila) | Source | Down |  |
| **HSP90AA1** | Heat shock protein 90kDa alpha family class A member 1 | Source | Up | Associated with virus packaging (68). |
| **HSP90AB1** | Heat shock protein 90kDa alpha family class B member 1 | Source | Up | Associated with virus packaging (68). |
| **HSP90B1** | Heat shock protein 90kDa beta family member 1 | Source | Up | Associated with virus packaging (68). |
| **IL11** | Interleukin 11 | Source | Up |  |
| **IL3** | Interleukin 3 | Source | Up |  |
| **JAG1** | Jagged 1 | Source | Up |  |
| **LPAR1** | Lysophosphatidic acid receptor 1 | Source | Down |  |
| **LPAR2** | Lysophosphatidic acid receptor 2 | Source | Up |  |
| **LPAR4** | Lysophosphatidic acid receptor 4 | Source | Down |  |
| **MAML1** | Mastermind like transcriptional coactivator 1 | Source | Down |  |
| **MAML2** | Mastermind like transcriptional coactivator 2 | Source | Down |  |
| **ANGPTL4** | Angiopoietin like 4 | Sink | Down |  |
| **BCL2L1** | BCL2-like 1 | Sink | Down |  |
| **IL17F** | Interleukin 17F | Sink | Down |  |
| **MCL1** | Myeloid cell leukemia 1 | Sink | Up |  |
| **PCK1** | Phosphoenolpyruvate carboxykinase 1 (soluble) | Sink | Up |  |
| **IL21** | Interleukin 21 | 1.62 | Down | IL-21 is critical for the development of optimal vaccine-induced primary but not secondary antibody responses against RABV infections (Dorfmeier et al., 2013) |
| **IFNB1** | Interferon, beta 1, fibroblast | 1.19 | Up | RABV P protein binds and inhibit Binding to IRF3 (Brzozka et al., 2006) |
| **EP300** | E1A binding protein p300 | 0.70 | Down |  |
| **TLR4** | Toll-like receptor 4 | 0.64 | Down | No sign of phenotype due to lacking TLR4. |
| **IL6** | Interleukin 6 | 0.58 | Up | Overexpression during infection (58,59), correlation among the IL-6 genes and the way of behavioral lateralization (60), involved in RABV pathogenesis (61). |
| **MYD88** | Myeloid differentiation primary response 88 | 0.44 | Down | Weakened RABV intervenes deadly disease while no MyD88 present, genetic adjuvanting with Myd88 improved the RVNA responses of a plasmid DNA rabies vaccine (90,91) |
| **NFKB1** | Nuclear factor of kappa light polypeptide gene enhancer in B-cells 1 | 0.13 | Down |  |
| **STAT3** | Signal transducer and activator of transcription 3 (acute-phase response factor) | 0.12 | Down | inhibits STAT3 nuclear accumulation (Lieu et al., 2013) |
| **TRAF6** | TNF receptor associated factor 6 | 0.08 | Up |  |
| **F2R** | Coagulation factor II (thrombin) receptor | 0.05 | Up |  |
| **CASP8** | Caspase 8, apoptosis-related cysteine peptidase | 0.00 | Up | Activation in RABV (Sarmiento et al., 2006) |
| **GNAI2** | Guanine nucleotide binding protein (G protein), alpha inhibiting activity polypeptide 2 | 0.00 | Down |  |
| **RBL1** | Retinoblastoma-like 1 | 0.00 | Down |  |
| **TFDP2** | Transcription factor Dp-2 (E2F dimerization partner 2) | 0.00 | Down |  |
| **STAT1** | Signal transducer and activator of transcription 1 | -0.01 | Down | RABV P protein binds and inhibit dimerization of STAT (Vidy et al., 2005; Brzozka et al., 2006), inhibit translocation (Brzozka et al., 2006; Moseley et al., 2009) |
| **AKT2** | v-AKT murine thymoma viral oncogene homolog 2 | -0.07 | Up | Hyper-phosphorylation of RABV P protein (Sun et al., 2008) |
| **AKT3** | v-AKT murine thymoma viral oncogene homolog 3 | -0.07 | Up | Hyper-phosphorylation of RABV P protein (Sun et al., 2008) |
| **JUN** | Jun proto-oncogene | -0.32 | Up | Activated in Rabies (Nakamichi et al., 2005) |
| **FASLG** | Fas ligand (TNF superfamily, member 6) | -0.48 | Up | Immune disruptive strategy of RABV to bring about apoptosis in T cell by overexpression in neuron (93). |
| **IRF7** | Interferon regulatory factor 7 | -0.60 | Up | RABV P protein averts tis activation (70) |
| **ADCY6** | Adenylate cyclase 6 | -0.70 | Up | the signal pathway from the stimulating regulatory component of the adenylate cyclase system to the unchanged activity of the catalytic subunit is defective (Koschel and Halbach, 1979; Koschel and Munzel, 1984) |
| **ADCY1** | Adenylate cyclase 1 (brain) | -0.73 | Down | the signal pathway from the stimulating regulatory component of the adenylate cyclase system to the unchanged activity of the catalytic subunit is defective (Koschel and Halbach, 1979; Koschel and Munzel, 1984) |
| **ADCY7** | Adenylate cyclase 7 | -0.73 | Up | the signal pathway from the stimulating regulatory component of the adenylate cyclase system to the unchanged activity of the catalytic subunit is defective (Koschel and Halbach, 1979; Koschel and Munzel, 1984) |
| **ADCY9** | Adenylate cyclase 9 | -0.73 | Down | the signal pathway from the stimulating regulatory component of the adenylate cyclase system to the unchanged activity of the catalytic subunit is defective (Koschel and Halbach, 1979; Koschel and Munzel, 1984) |
| **IRF3** | Interferon regulatory factor 3 | -0.78 | Up | RABV P protein binds and avert binding to IFNB1 (53), inhibit IRF3 phosphorylation (70,96) |
| **JAK2** | Janus kinase 2 | -0.79 | Up |  |

#### Interferon circumvent

To escape innate and adaptive immunity, rabies perturbs Jak-Stat signaling by influencing interactions and expression. As shown in figure 5A, two modules of RISN are likely to interfere with this signaling pathway. The inhibition of dimerization of Stat proteins and accumulation in nucleus by viral P protein are previously described (Vidy et al., 2005; Brzozka et al., 2006; Moseley et al., 2009; Lieu et al., 2013). Our findings showed the down-regulation of STAT proteins, which is plausible considering the feedback self-loop control on these proteins. This finding is supported by up-regulation of the feedback inhibitor known as the SOCS protein. The high betweenness value of JAK2 and its up-regulation indicate the role of activating innate and adaptive defense systems of infected cells against rabies. Additionally, the low IRI value of JAK2 and transfer of information towards the “toward proliferation and survival” modules highlight its importance in rabies etiology. The expression alterations of IL6 and IL21 were also reported in rabies infection previously (Hemachudha et al., 1993; Megid et al., 2006; Quaranta et al., 2008; Dorfmeier et al., 2013; Srithayakumar et al., 2014). The down-regulation of IL21 and its receptors, IL21R, IL15R and IL17F, following the down-regulation of STAT3, is very significant in the observed dampened immune system of the rabid given their role in proliferation and maturation of natural killer cells. Unlikely, the up-regulation of IL6, a neuroprotective cytokine, reinforces the anti-apoptotic effects of the rabies wild type strain. Both of these play a propagation role in this network with IRI values above one. Despite vastly interfering with the Jak-Stat signaling, rabies infection could not decrease the expression of the famous antiviral molecule, IFNB1, and cell could have upregulated it against the infection. Surprisingly, however, the virus chooses another strategy to skip the interferon mechanism (Faul et al., 2010). This alternative plan is to perturb IFNB1 activation via IRF3. The over-expression of RORA and RORC is also important since they affect the circadian rhythm, calcium-mediated signal transduction and anti-inflammatory responses. It seems that the up-regulated MCL1 and BCL2L1, act in favor of survival, inflammation attenuation and apoptosis inhibition of neurons and disrupt endocytic vesicle retrieval. Also, decrease in function of CIITA and TBX1 causes decrease in the function and efficiency of TH1 and TH2. Overall under-expression of the Notch signaling pathway, including DLL1, NOTCH1 and RBPJ, along with the up-regulation of JAG1, an inhibitor of NOTCH1, is indicative of malfunction in cell-cell communication in CNS and neuronal self-renewal mechanism.

#### Toward proliferation and survival

Similar to other viral infection, the RABV hijacks the proliferation machine of cells to generate virions as much as possible. To achieve this goal, the strategy of the virus is to keep the cell alive and active and amplifies the production rate. Rabies achieves this by using cell envelope and preventing apoptosis. In fact, there was an inverse correlation between induction of apoptosis and the potency of a virus strain to invade the brain. This suggests that suppression of apoptosis may well be a strategy for neuro-invasiveness of pathogenic RABV and progression through the nervous system (Thoulouze et al., 2003a; Thoulouze et al., 2003b; Larrous et al., 2010). As it is shown in figure 5B, up-regulated AKT2 and AKT3 in the tyrosine kinase module play a central role in WHAT??? It has been previously reported that AKT signaling is hijacked by non-segmented RNA viruses such as vesicular stomatitis virus (VSV) via phosphorylating P proteins (Sun et al., 2008). It has also been suggested to use AKT inhibitors as an anti-RABVdrug. Our findings underscore the importance of this signaling pathway in neuronal cells where AKT signaling is not normally hyperactive. This activated pathway, prompted by G viral proteins, may result in the activation of proliferation and growth machinery, and help viral protein folding and packaging via over-expression of HSPs (Lahaye et al., 2009; Lahaye et al., 2012). TSC2 also triggers apoptosis in immune cells via high representation of FASLG. These results in parallel with those in Sun et al. (REF) highlights the need for studying AKTs and anti-AKTs in rabies models.

Other Serine or Tyrosine kinases, including ITK, IKBKB and RAF1 and PIK3CG were under-expressed, thus resulting in the dampening of the inflammatory response especially with the up-regulation of anti-inflammatory proteins such as CHUK and cell proliferatory proteins, NRAS and KRAS. IRF3, IRF7 and IFNB1, which are upregulated naturally in response to viral infection, could not stimulate an immune response due to the activation of the P viral protein. Besides, IRF3 and IRF7 are involved in AKT activation and transformation of inflammatory to anti-inflammatory macrophages (Rieder et al., 2011; Tarassishin et al., 2011). The over-expressed AKT genes also inactivate FOXO3 and therefore disrupt the cell efforts toward apoptosis (Tarassishin et al., 2011). Expression of ANGPTL4 that causes Anoikis, a type of programmed cell death, is also decreased after this module activity (Terada and Nwariaku, 2011). The expression alteration of proteins involved in cell cycle regulation including RB1, RBL1, RBL2, TFDP2 and E2F1 is indicative of the triggering disruption and hijacking by the virus.

#### Neuropathological clue

Hitherto, the underlying mechanism of escape from the immune system, apoptosis prevention and virus production in rabies infection is demonstrable by these modules (Fig 5C). However, importantly, the main cause of death in rabid is cardiac arrhythmia and breathing pattern disorders, for which its molecular basis should be tracked elsewhere. Meanwhile, trace of chemokines such as CCL3, CCL5 (a neuron survival factor) and CXCL10 in rabies infection has been previously detected (Nakamichi et al., 2005; Johnson et al., 2008; Li et al., 2012; Huang et al., 2014). These molecules along with their receptors are involved in the blood brain barrier (BBB) permeability and recruitment of different T cells (Fig. 5C). In the natural cell cycle, MYD88 expression leads to an increase in expression of NFᴋB and hence programmed cell death. Seemingly, the expression and function of of MYD88 in addition to NFᴋB is decreased (Fig. 5C) which can lead to the suppression of apoptosis. Diverse chemokines as source nodes in RISN and information transduction to adenylate cyclases and MAP kinases via GNAI2 is likely to be a critical clue to discovering the etiological mechanism of rabies fatality.

Adenylyl cyclases (ADCYs) are central components of signaling cascades downstream of many G proteins. In the mammalians, of the ten ADCY isoforms identified, nine (ADCY1-9) are transmembrane proteins, whereas ADCY10 is a soluble isoform that lacks the transmembrane domains (Sunahara et al., 1996; Conley et al., 2013; Birrell et al., 2015). Although in the RISN the expression of *ADCY1* and *ADCY9* was decreased, the level of ADCY6 and ADCY7 was increased, the overall outcome is probably reduction of signal flow rousted by GPCR and ADCY. All ADCY isoforms catalyze the conversion of ATP to cyclic AMP (cAMP) and pyrophosphate. cAMP is a messenger involved in many biological processes including cell growth and differentiation, transcriptional regulation, apoptosis and various other cellular functions (Patel et al., 2001). The main protein kinase activated by cAMP is protein kinase A (PKA). PKA transfers phosphate groups form ATP to proteins including ion channels on the cell membrane. Similar to changes in enzyme activity following biochemical modification, phosphorylation of ion channel proteins may also cause conformational changes and consequently increase chances of channel opening leading to depolarization of postsynaptic neurons, resulting in firing an action potential and altered electrical activity properties. On the other hand, ADCYs are integrated in lipid rafts and caveolae, and implicated in local cAMP micro-domains in the membrane (Schwencke et al., 1999). Subcellular compartmentalization of protein kinases (such as PKA) and phosphatases, through their interaction with A kinase anchoring protein (AKAPs), provides a mechanism to control signal transduction events at specific sites within the cell (McConnachie et al., 2006). The RABV may interfere with the lipid raft and the micro-compartment associated with the cAMP-AKAP-PKA complex and thus alter ion channel function, eventually leading to neuronal dysfunction. In line with this, it has been reported that NMDA and AMPA glutamate receptors form complexes with cytoskeletal and scaffold proteins in the post-synaptic density (PSD) (Kennedy, 1997; Ziff, 1997). Interestingly, AKAP binds to PSD in complexes with NMDA and AMPA receptors (Colledge et al., 2000). It is also thought that regulation of this molecular architecture is essential for controlling glutamate receptors in hippocampal long-term potentiation (LTP) and long-term depression (LTD) synaptic plasticity (Luscher et al., 2000; Tomita et al., 2001). Our data showed that the expression of PRKACA and AKAP13 subsequent to GNAI2 decreased significantly (Supplementary File 7). It therefore seems that these complexes may be a preferential target of viruses to hijack cellular machinery.

Neuropathological observations indicate that functional alterations precede neuronal death, which is responsible for the clinical manifestation and fatal outcome in rabies.. Indeed, Gourmelon et al. reported that disappearance of rapid eye movement (REM) sleep and the development of pseudoperiodic facial myoclonus are the first manifestations in the EEG recordings of mice infected with the challenge virus standard (CVS) of fixed RV (Gourmelon et al., 1986). It has also been reported that electrical activity of brain terminates thirty minutes prior to the cardiac arrest, indicating that cerebral death occurs before vegetative function failure in experimental rabies (Fu and Jackson, 2005). Considering the increased activity of voltage-gated channels by phosphorylation in response to PKA stimulation, initiating a signaling pathway from ADCY to ion channel functioning could be a possible mechanism by which the RABV hijacks the neurons. Consistently, Iwata et al. showed that ion channel dysfunction occurs in mouse neuroblastoma cells infected by RV (Iwata et al., 1999). They reported that not only the functional activity of voltage-dependent sodium and inward rectifier potassium channels were decreased, but also the resting membrane potential was decreased, indicating membrane depolarization. Therefore, decreased activity of these channels could preclude infected neurons from firing action potentials and generating synaptic potentials,, and may lead to functional impairment. Fu et al. observed that neurotransmitter releases from rat hippocampus, after inoculation with RV CVS-24, was increased at day 1, reached a peak at day 3 and then declined by day 5. Manifestations of clinical signs of rabies were consistent with day 5 of inoculation when neurotransmitter release was equal or below the level prior to infection, suggesting that neurons are no longer capable of releasing neurotransmitters at the synaptic junctions and this may be the underlying basis of clinical signs including paralysis (Fu and Jackson, 2005).

Since there is a paucity of data pertaining to rabies influence on physiological processes, particularly on neuronal electrophysiological properties, further studies need to be undertaken to confirm whether neuronal dysfunction occurring in rabies infection is due to an aberrant signaling pathway initiating from ADCY-cAMP-PKA and finalizing with ion channel phosphorylation. It should be noted that any alterations in different ion channels may result in dysfunction of neurons and brain regions which are responsible for vital tasks including attention, thinking and respiration.

### Manually curated version of RISN

To simplify RISN, signaling pathways were manually extracted and merged based on the currently available data in KEGG, including WNT, MAPK/ERK, RAS, PI3K/AKT, Toll-like receptor, JAK/STAT and NOTCH signaling pathways. The information flow from diverse ligands to various transcription factors is illustrated along with differential expression. As shown in Fig 6, information is converged toward several important proteins including PLC, MAPK1/2, PIK3, PKC, and JAK, and is then diverged toward several distinct transcription factors and finally end-point biological processes.

**Fig 6.**
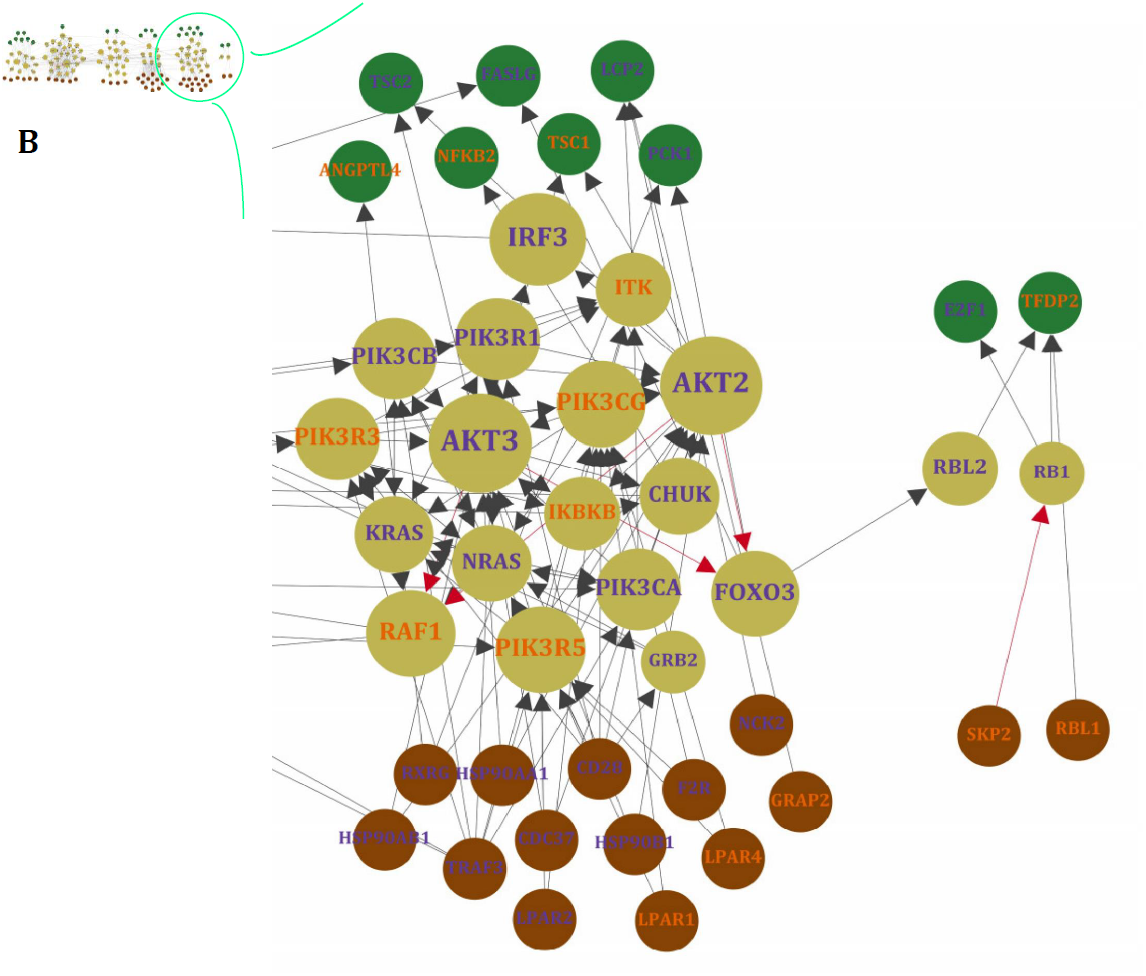
The manually curated version of RISN. The over-and under-expressed genes are colored green and yellow, respectively. The cellular membrane and nuclear membrane are depicted by gray solid and dashed lines, respectively. Viral proteins and RNA are depicted by non-circular and rectangular objects, and DNA and small metabolite molecules are depicted by circular shapes. The phosphorylated P proteinactivated by AKT is denoted by P*, however, for the sake of simplicity, its process is not shown in the signaling pathway. This symbol with different caps was used to show activatory and inhibitory effects of RABV on RISN.

Our analysis revealed that two of three WNT signaling pathways were altered in rabies infected cells. The canonical WNT pathway (WNT/β-catenin) along with the noncanonical planar cell polarity (PCP) pathway were apparently active in infected neurons but the noncanonical WNT/calcium pathway was not induced. The PCP pathway is involved in up-regulation of components of the downstream pathway and cytoskeletal rearrangements of which the latter may implicate this pathway in cytoskeletal changes in neurons. This is consistent with previous studies reporting cooperative cytoskeletal changes (restructuration) for viral protein transportation and viral localization (Sagara et al., 1995; Ceccaldi et al., 1997; Song et al., 2013; Zandi et al., 2013).

There is also evidence of crosstalk between WNT and MAPK/ERK signaling pathways. It seems that in the rabid brains the MAPK/ERK signaling pathway, via cAMP-PCREB signaling, is involved in neuromelanin biosynthesis of which its accumulation depletes iron ions as observed in some neurodegenerative diseases such as Parkinson’s disease (Good et al., 1992). Iron deficiency may also contribute to defective dopaminergic interaction with neurotransmission systems (Youdim, 2008). This is, however, a speculation and needs experimental validation in rabies infection cases.

Additionally, RAS signaling is activated through the C-Kit receptor and diverge toward PIK3 and MAPK/ERK signaling pathways. Downstream of RAS activation (ERK signaling and AKT) is highly complex but generally contributes to cell growth (Bender et al., 2015). Activated RAS signaling suppresses PKR-mediated responses to interferon response and double-stranded RNA degradation. Normally, viral transcripts trigger PKR phosphorylation and activation, and finally inhibit infection. Therefore, the RABV may replicate silently in RAS activated cells (Mundschau and Faller, 1994; Russell, 2002). This data-based hypothesis also requires experimental validation in rabid cases.

The AKT signaling pathway plays a critical role in the replication of the RABV similar to other non-segmented negative-stranded RNA viruses. Heavy phosphorylation of viral proteins (P protein) is mainly mediated via AKT activity (Sun et al., 2008). Subsequently, the activated P protein plays a crucial role in other signaling pathways such as Toll-like receptor and JAK/STAT signaling pathways which are responsible for viral genome detection and immune-modulatory functions against rabies, respectively. Accordingly, the viral G protein activates AKT signaling through phosphorylation and localization of PTEN (Terrien et al., 2012). The consequences of the activation and crosstalk of these signaling pathways are reduced apoptosis, cell survival and blocked cell cycle progression. Neuronal dysfunction, inhibition of apoptosis and limitation of inflammation have been previously stressed by Gomme *et al*. (Gomme et al., 2012a). It seems that these processes have been evolutionarily acquired to complete virus lifecycle and transfer to the new host. They also showed that most of DEGs are involved in signaling transduction and nervous system function, and therefore affect cell behavior by decreasing neurite growth, organization of cytoskeleton and cytoplasm, and microtubule dynamics.

It has been demonstrated that rabies infection up-regulates expression of CXCL10 and CCL5 proteins in a ERK1/2-, p38- and NFkB-dependent manner (Nakamichi et al., 2004; Nakamichi et al., 2005). CXCL10 is a major chemo-attractant of Th-1 cells. The up-regulation of interferon, chemokines, interleukin (IL) and IL-related genes were previously observed by Sugiura *et al*. (Sugiura et al., 2011). They also reported the signaling pathways involved in rabies infection including interferon signaling, IL-15 production and signaling, and Granzyme B signaling which trigger apoptosis in immunetarget cells.

ERK and p38MAPK along with BCL2 family and the FasL receptor are important apoptosis committers. Currently, it is known that RISN inhibits apoptosis and also suppresses cell proliferation (Gomme et al., 2012a). Also, JAK/STAT signaling is activated through its receptors, but as mentioned earlier STAT dimerization is inhibited via viral P protein activity. Therefore, downstream signaling of STATs, which is critical for interferon signaling and viral defense, is suppressed. Concomitant activation of JAK/STAT and AKT signaling pathways has a pro-survival function in neurons (Junyent et al., 2010). In a time-course study, Zhao *et al*. studied the gene expression profile of infected microglial cells and indicated some affected signaling pathways at different time points (Zhao et al., 2013). The MAPK, chemokine, and JAK-STAT signaling were also shown to be implicated in rabies infection along with other innate and adaptive immune response pathways. These pathways were also detected in two other independent studies on CNS of infected mice (Zhao et al., 2011; Zhao et al., 2012b).

Viral pattern recognition is critical for early innate immunity response and modulation of pathogen-specific adaptive immunity. TLR3, a member of the TLR family which are pattern recognition receptors in cells, is increased in the cytoplasm of rabies infected cells. It plays major functions in spatial arrangements of infected cells and viral replication, and is observed in endosomes and Negri bodies which are only formed in the presence of TLR3 (Ménager et al., 2009).

NOTCH signaling is important for cell communication, neuronal function and development in spatial learning and memory (Costa et al., 2003). Our data indicate that this signaling pathway is active in rabid brains. More detailed examination of the role of this signaling pathway in rabies infection is warranted.

### Experimental validation of expression alterations

Gene expression analysis of a number of randomly selected DEGs has been performed previously based on RT-qPCR as a routine validation method of microarray-based expression profiles (Zhao et al., 2011; Zhao et al., 2012a; Zhao et al., 2012b; Zhao et al., 2013) or fluorescent bead immunoassay (Sugiura et al., 2011). In all cases, the former experimentally confirmed DEGs identified in SHIDEGs were *STAT1, STAT3, SOCS2, IRF1, IRF3, IRF7, IFNAR2, SH2D1A, CCL3, CCL5, CCRL2, CXCL10, Mx1, IFIT3, OASL2, USP18, IL6, IL10, IL23A* and *RTP4*. However, we examined another independent random gene set among RISN genes as a further step of validating the microarray-based results. The differential expression of *AKT3, GNAI2* and *IL21* was analyzed by comparing expression levels in murine neuroblastoma cells infected by the wild type RABV with control uninfected cells using RT-qPCR. The results indicated that the direction of differential expression of all three genes were consistent between RT-qPCR results and data integrated from multiple microarray chips (Fig 7, supplementary file 8 and 9). These results confirmed the down-regulation of *GNAI2* and *IL21*. AKT values and statistical tests states that the expression of AKT is not up regulated in infected samples. Our findings based on delta Ct method and comparing the raw values of AKT gene at both samples with the referenced values; however, firmly confirm that the value of gene is indeed up regulated in infected samples.

**Fig 7.**
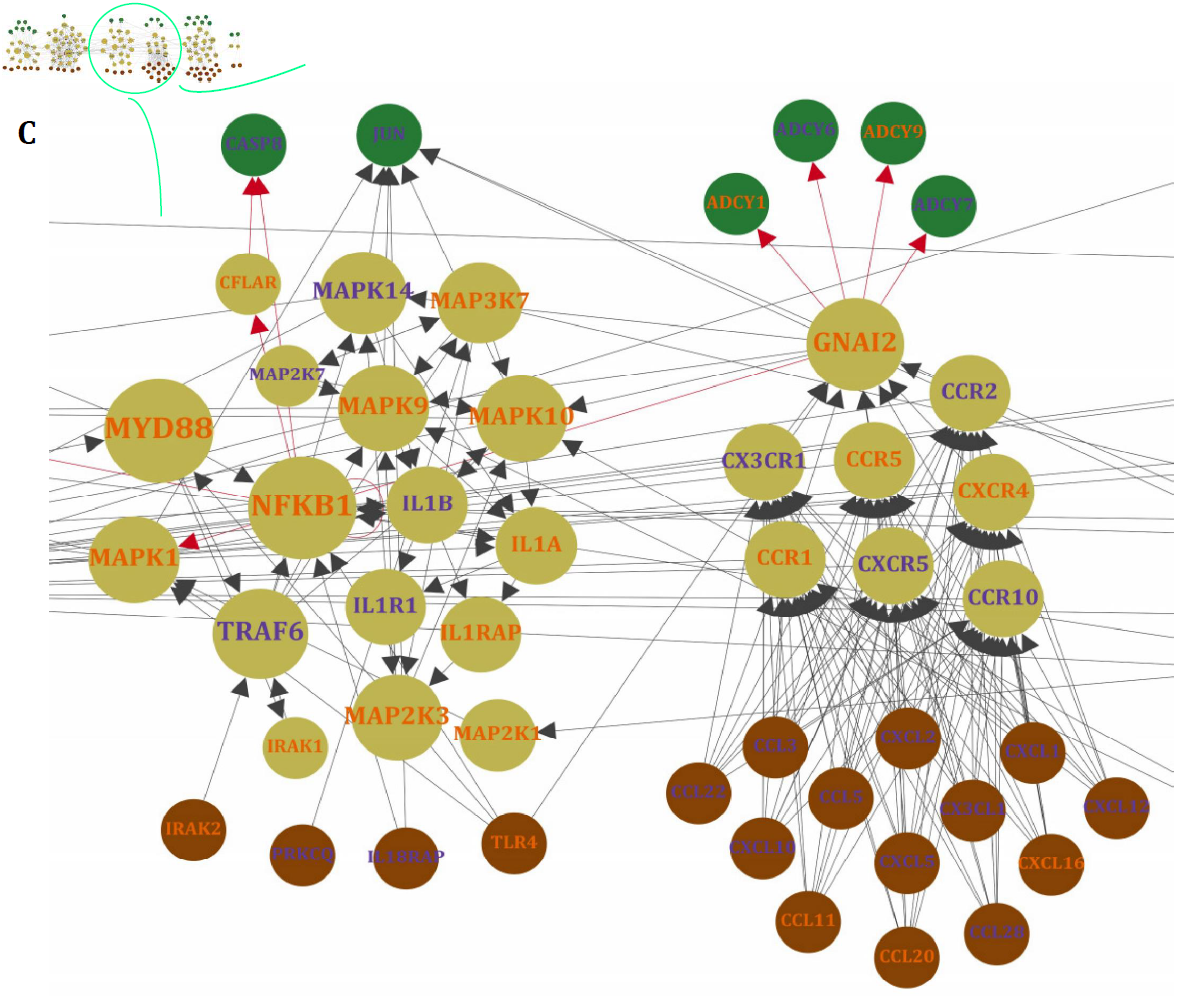
Experimental validation of microarray-based expression results. A) Mock infected N2a cell culture image showing a normal morphology. B) N2a cells infected with CVS strain of RABV with multiplicity of infection (MOI=3), stained by FITC conjugated anti-rabies nucleocapsid polycolonal antibody. The images were captured 24h post infection. C) Expression profile of three randomly selected genes acting in different signaling pathways of the rabies-implicated signaling network (RISN). Expression levels were quantified using RT-qPCR in triplicates. The error bar plots indicate Mean +/-SD and include the corresponding p-value of the statistical significance test. The vertical axis measures the negative inverse value of the logarithm of the mean value for each replicate using the delta Ct method.

## Discussion

Rabies is a fatal neuropathological disorder. The fatality of this infection is not because of neurological damage or neurohistopathological signs, but due to neurophysiological disruption of vital signs such as regular heart beat and respiratory rhythm. Other evidences that highlight this neural malfunction are known rabies symptoms such as hydrophobia, photophobia, and paralysis of facial and throat muscles. Although rabies infected cells can mount an innate immune response against this infection, the virus can control the expression and function of the proteins involved in the induction of apoptosis and efficiently suppresses the antiviral innate immune response. From a pathobiological point of view, we acknowledge that the RISN and the previously reported pathways which lead to the spread of RABV can also be triggered by unrelated viruses including other neurotropic RNA viruses, measles, and influenza. This infection would yield similar gene expression profiles by activating general host responses including activation of stress response, innate immune response and interferon signaling signatures.

Knowledge-driven studies are mostly non-automatic, heuristic, expert-dependent and evidence-based surveys. Although this strategy of problem solving is valuable in identification of novel findings, it suffers from certain limitations and subjective biases. With the emergence of omics technologies, data-driven studies have exploited the large and ever-growing publicly available deposited datasets as a complementary approach to knowledge-based studies (Sun et al., 2012). Data-driven approaches are computationally demanding and require complex interpretations but this is dependent directly to the original data itself (Hua et al., 2006; Sun et al., 2012). Here, we combined data- and knowledge-driven studies to potentially identify a less-biased signaling network of rabies infection. We thus undertook a systematic approach which initiated with a data-driven approach and was then extended by a comprehensive complementary knowledge-based approach. In addition, we included all available high-throughput whole-transcriptome datasets in a horizontal (meta-analysis) and super-horizontal (miRNA and mRNA) integration approach. Critically, signaling pathways were then used as a scaffold for data integration to identify key players in signaling pathways and genes. Finally, we constructed a bird’s-eye view map, RISN, of signaling deviations including host-pathogen interaction data. Uniquely, this signaling pathway illustrates the host-rabies interaction signature.

Our results indicate that seven signaling pathways including 1) WNT, 2) MAPK/ERK, 3) RAS, 4) PI3K/AKT, 5) Toll-like receptor, 6) JAK/STAT and 7) NOTCH are involved in controlling cell cycle, cell survival, viral replication and folding, synapse regulation and regulation of immunity. Among the many involved proteins, divergence and convergence of signals indicates that PLC, MAPK1/2, PIK3, PKC and JAK are potentially the most critical of all in rabies pathogenesis. Interestingly, signals are converged toward these proteins and are then diverged toward several distinct transcription factors and end-point biological processes (Fig. 8).

**Fig 8.**
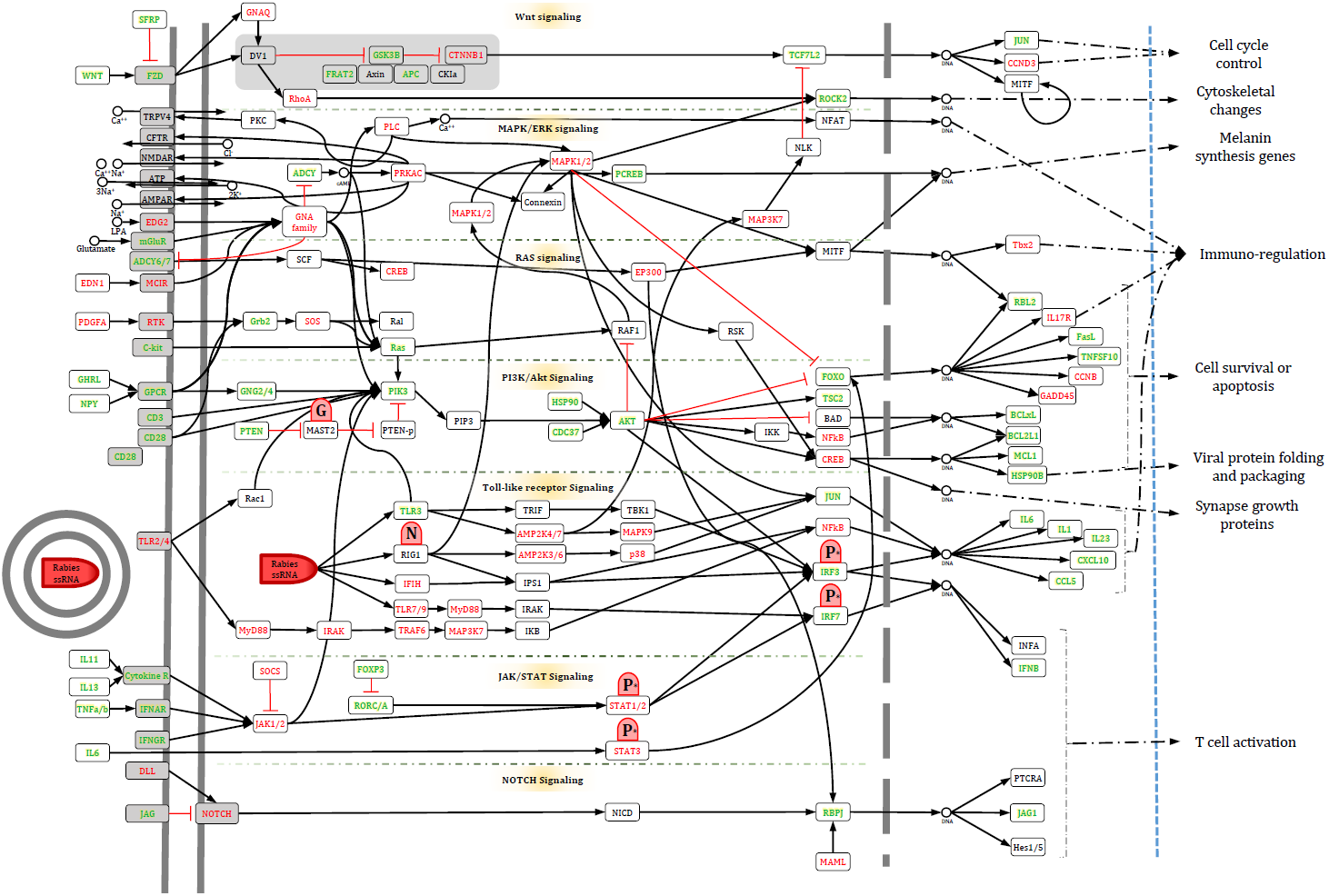
The manually curated version of RISN in a nutshell. To summarize the biological processes involved in the infected neuron, the signaling pathways along with their triggers and consequences are delineated based on Figure 6. Activated and inhibited pathways are colored green and red, respectively. The different caps of this symbol were used to show activatory and inhibitory effects of RABV on RISN.

In addition to confirming former reports on the inhibition of apoptosis in neurons, RISN provided molecular evidence of interferon escape and neural cell death prevention in rabies infection. This finding is significant given that it explicates how the virus continues to parasitically multiply without any neural host cell damage. Data herein suggest that, the RABV hijacks the phosphorylation machinery of the cell to facilitate its own replication. Also, the tight regulation of recruited immune cells by the virus is demonstrated. The network analysis also shed light on the gene set central to rabies infection, all of which were bottlenecks in RISN. Moreover, based on RISN, we hypothesize that modifying certain signal transduction apparatus involved in rabies etiology such as the cAMP or AKT signaling pathway may instigate an effective immune response which will consequently diminish the fatality of the rabies infection.

The systems biomedicine approach employed in this study provided a better understanding of the underlying signaling network of this infectious disease. Further independent validation of the RISN potentially provides a molecular framework for intervention and development of novel effective treatments for the late stages of this neglected disease.

## Acknowledgement

The authors wish to thank Karl-Klaus Conzelmann, Monique Lafon and Hervé Bourhy for providing us with in-depth knowledge on rabies and their critical evaluation of the manuscript.

## Supporting information Captions

**Supplementary File 1:** The selected datasets for rabies data integration.

**Supplementary File 2:** The computational scripts plus all raw data used in this study

**Supplementary File 3:** The degree distribution of the refined SHIDEG-PPIN.

**Supplementary File 4:** The gene ontology enrichment analysis of differentially expressed and central (degree and betweenness) genes in SHIDEG-PPIN.

**Supplementary File 5:** The full results of pathway enrichment analysis of SHIDEG-PPIN modules by ClueGO.

**Supplementary File 6:** The KEGG enriched pathway similarity matrix and the module interconnectivity matrix of RISN.

**Supplementary File 7:** Properties of RISN nodes and list of all edges.

**Supplementary File 8:** The over-and under-expression of 694 DEGs in the unrefined RISN.

**Supplementary File 9:** Properties of refined SHIDEG-PPIN nodes and list of edges.

---

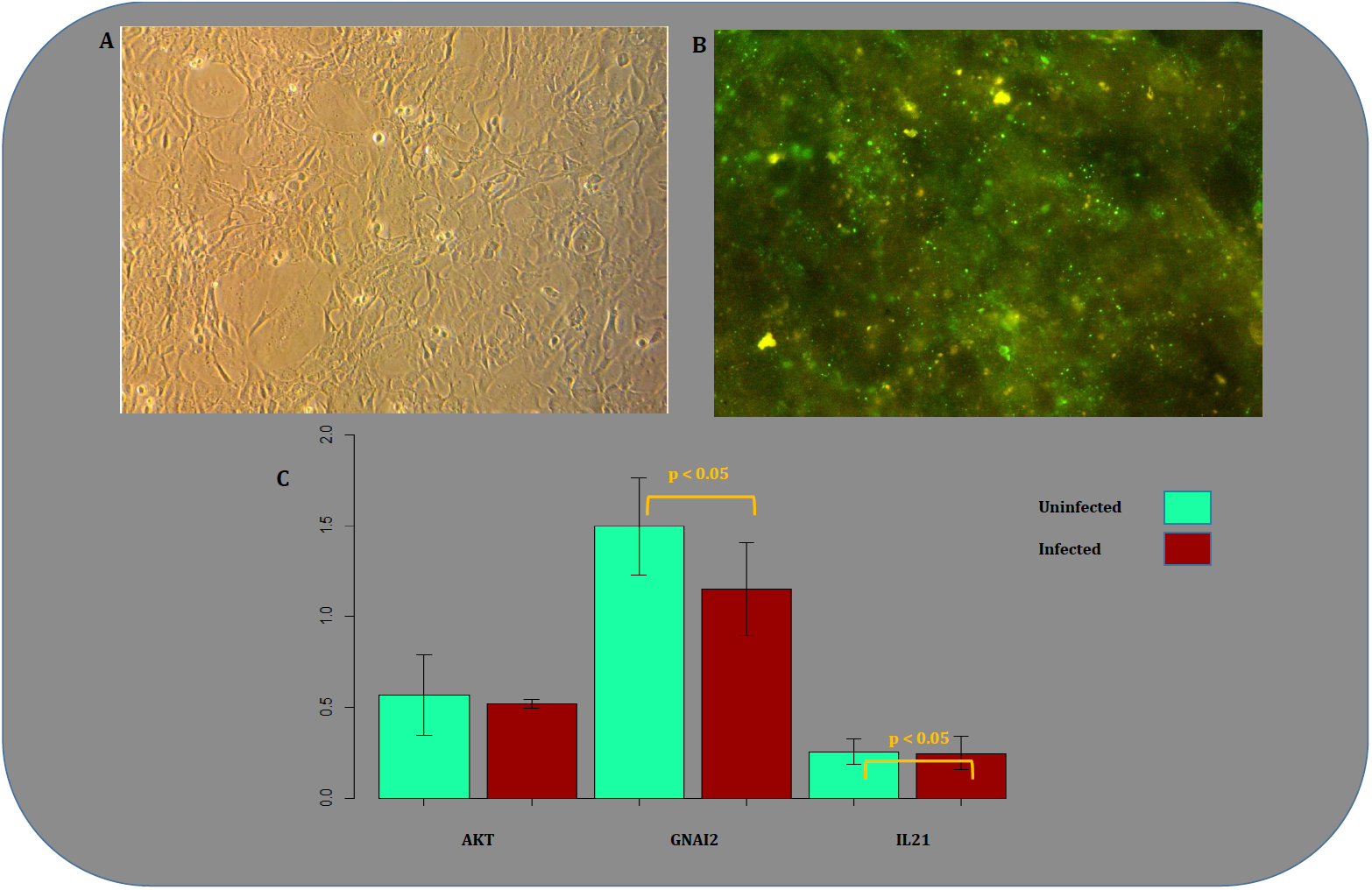

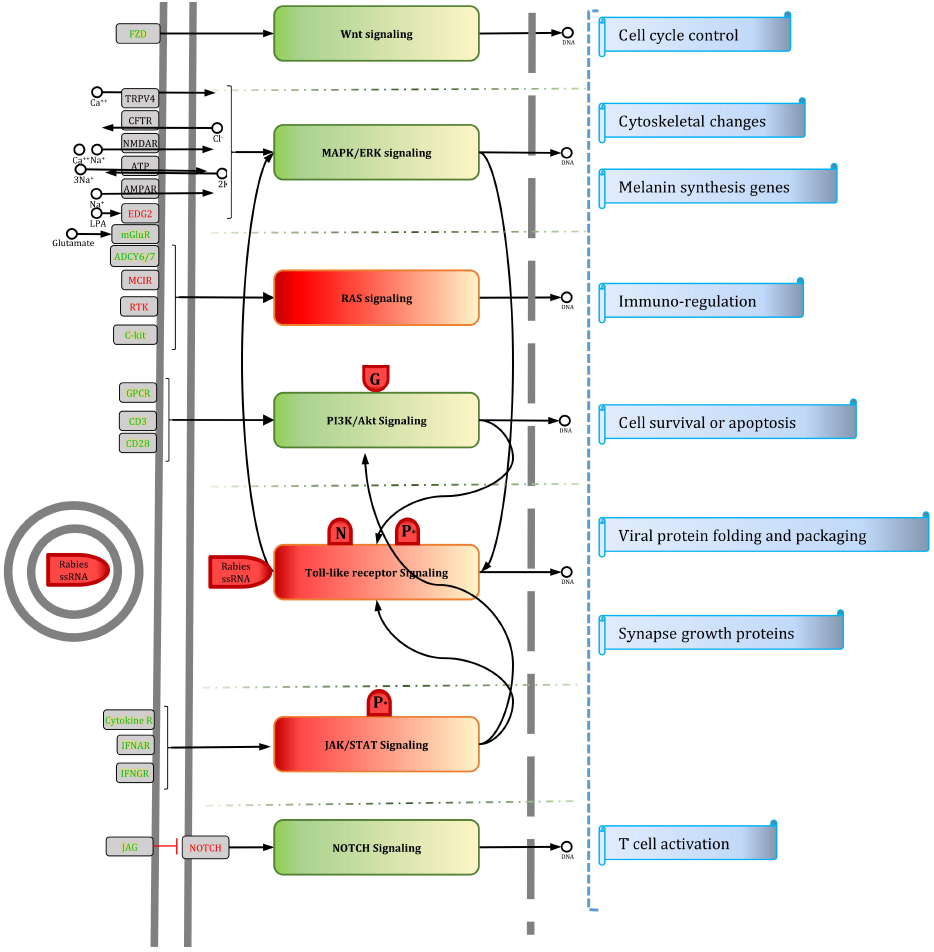

